# Improved methods and optimized design for CRISPR Cas9 and Cas12a homology-directed repair

**DOI:** 10.1101/2021.04.07.438685

**Authors:** Mollie S. Schubert, Bernice Thommandru, Jessica Woodley, Rolf Turk, Shuqi Yan, Gavin Kurgan, Matthew S. McNeill, Garrett R. Rettig

**Affiliations:** Integrated DNA Technologies, Inc., Coralville, IA, 52241, USA; BGI Genomics, BGI-Shenzhen, Shenzhen 518083, China

## Abstract

CRISPR-Cas proteins are used to introduce double-stranded breaks (DSBs) at targeted genomic loci. DSBs are repaired by endogenous cellular pathways such as non-homologous end joining (NHEJ) and homology-directed repair (HDR). Providing a DNA template during repair allows for precise introduction of a desired mutation via the HDR pathway. However, rates of repair by HDR are often slow compared to the more rapid but less accurate NHEJ-mediated repair. Here, we describe comprehensive design considerations and optimized methods for highly efficient HDR using single-stranded oligodeoxynucleotide (ssODN) donor templates for several CRISPR-Cas systems including *S.p.* Cas9, *S.p.* Cas9 D10A nickase, and *A.s.* Cas12a delivered as ribonucleoprotein complexes with synthetic guide RNAs. Features relating to guide RNA selection, donor strand preference, and incorporation of blocking mutations in the donor template to prevent re-cleavage were investigated and were implemented in a novel online tool for HDR donor template design. Additionally, we employ chemically modified HDR donor templates in combination with a small molecule to boost HDR efficiency up to 10-fold. These findings allow for high frequencies of precise repair utilizing HDR in multiple mammalian cell lines. Tool availability: www.idtdna.com/HDR

## INTRODUCTION

CRISPR-Cas systems have revolutionized genomics by enabling efficient and precise genome editing in a wide variety of biological systems, including eukaryotic cells.^1–5^ These systems require an RNA-guided DNA endonuclease and a target-specific guide RNA (gRNA) to generate a double-stranded break (DSB) at a desired genomic location, which must be flanked by a short protospacer adjacent motif (PAM). *Streptococcus pyogenes* Cas9 (*S.p.* Cas9) is one of the most commonly used CRISPR enzymes for genome editing. The native gRNA for Cas9 is hybridized from two RNA molecules: a CRISPR RNA (crRNA) and a universal, trans-activating crRNA (tracrRNA).^6^ The two strands of the gRNA can also be combined as a single unimolecular structure to form a single-guide RNA (sgRNA).^7^ Association of Cas9 protein with a gRNA forms a ribonucleoprotein (RNP) complex, which surveys a dsDNA substrate and generates a DSB when its complementary target sequence with a PAM is recognized by an active Cas9 RNP complex.^8–10^

Recent reports have demonstrated that RNP delivery of nucleases has benefits over plasmid delivery as it enables a faster onset of action, reduces off-target cleavage, and eliminates the risk of random plasmid integration into the host genome.^9, 11, 12^ At the same time, RNP delivery allows for the use of chemically modified gRNA with improved stability and reduced toxicity.^13^ In addition, generating RNP complexes *in vitro* prior to delivery allows accurate control of the ratio of protein and gRNA to maximize RNP complexation efficiency. This also enables the formation of each gRNA:protein complex independently which mitigates the competition for Cas9 protein by other intracellular RNA molecules or by different gRNAs during multiplexing experiments.^14^

*S.p.* Cas9 contains two endonuclease domains (HNH and RuvC) that function together to generate a blunt DSB by each domain cleaving opposite DNA strands. Inactivating one of the two endonuclease domains results in Cas9 variants called “nickases”: the RuvC-inactive variant (Cas9 D10A) nicks the target (gRNA complementary) strand, while the HNH-inactive variant (Cas9 H840A) nicks the non-target (gRNA non-complementary) strand.^7^ Cas9 nickases can be used with an individual guide to induce single DNA nicks and induce a repair pathway termed alternative-HDR.^15, 16^ However, it is more common and often more efficient to perform genome editing at DSBs generated by using a nickase with a pair of gRNAs targeting opposite DNA strands in a “paired nicking strategy”.^17^ It has been demonstrated that nickases allow for the reduction of off-target editing by ∼50-1500 fold in comparison to Cas9 WT.^17–19^ At the same time, the paired nicking strategy can facilitate highly robust editing in many model systems, including mammalian tissue culture, mouse zygotes, plants, yeast, and bacteria.^1, 17–26^

Cas12a enzymes are also RNA-guided double-stranded DNA nucleases that provide an alternative to the commonly used *S.p.* Cas9 nuclease with similar editing outcomes. Unlike *S.p.* Cas9, which recognizes an NGG PAM sequence, *A.s.* Cas12a recognizes a TTTV (V = A/G/C) PAM site which allows for a broadened range of targeting sites in AT-rich regions. Cas12a relies on a single, short (41-44 nt) gRNA and generates staggered DSBs with 5’ overhangs.^4^ In addition, Cas12a has been shown to be advantageous due to intrinsically high specificity, reducing the potential for off-target cleavage.^27, 28^ However, Cas12a also has non-specific single-stranded DNase (ssDNase) activity that is activated upon binding to the target DNA strand.^29^ This could potentially impact the ability of Cas12a to mediate efficient HDR if the ssODN is degraded before it is able to act as a donor template.

To facilitate genome editing, CRISPR-Cas enzymes are used to generate a DSB at a genomic locus which can then be repaired by a variety of endogenous cellular repair pathways including non-homologous end joining (NHEJ) and homology-directed repair (HDR).^30, 31^ NHEJ is an imperfect process and commonly creates small insertions or deletions (indels), which can be exploited to introduce diverse, but reproducible genetic mutations or gene knockouts.^32^ On the other hand, HDR is a process that can lead to precise sequence alterations at specified genomic locations but requires the use of a carefully designed HDR donor template that contains sequences homologous to the specific sequence flanking the cut site, defined as ‘homology arms’. However, rates of repair by HDR are often slow compared to the more rapid but less accurate NHEJ-mediated repair.^33^ For small mutations or insertions, a ssODN can be used as the HDR donor template.^34–37^ These are readily available up to 200 nucleotides (nt) in length as chemically synthesized oligos, allowing for insertions up to 160 nt (maintaining, at minimum, 20-nt homology arms).

Two distinct pathways for the incorporation of single-stranded donor templates at a DSB have been proposed – single-strand DNA incorporation (ssDI) and synthesis-dependent strand annealing (SDSA), with SDSA being preferentially utilized as the repair path for ssODN donor templates in the presence of a DSB.^38^ Previous studies have examined design considerations for ssODNs when using CRISPR-Cas enzymes. The optimal length of homology arms has been reported to be as little as 30-nt in length on either side of the DSB, and it has been demonstrated that asymmetric donor oligos can improve HDR.^39, 40^ HDR efficiency is highest when the intended edit is placed near the DSB and is greatly reduced at loci distal to this event.^35, 41^ In addition, reports have indicated that there may be a preference for utilizing a donor oligo with sequences either complementary or non-complementary to the gRNA.^37, 42–44^ CRISPR-Cas9 can also re-cut dsDNA after a desired repair outcome if the protospacer and PAM sequence remains unaltered, lowering perfect HDR efficiency. This outcome can be prevented by strategically incorporating blocking mutations into the donor template.^35, 45^ Other approaches to improving HDR are of great interest to the genome editing community. It has been previously reported that incorporating chemical modifications such as phosphorothioate (PS) linkages may improve HDR when using ssODN donors.^39, 46^ Another route to improving HDR frequency is using chemical compounds that inhibit key DSB repair enzymes that play a role in the competing NHEJ pathway. Several chemical compounds have been reported to increase HDR.^47–49^ In this work, we thoroughly investigated design features for both *S.p.* Cas9 and *A.s.* Cas12a nucleases relating to gRNA selection, donor strand preference, the placement and composition of blocking mutations, and the number of blocking mutations that are required for maximum HDR efficiency. We additionally investigated alternate end-blocking oligo modifications to further stabilize the ssODN from exonuclease activity and have developed a novel modification, which is incorporated into Alt-R HDR Donor Oligos, that improves upon previously reported constructs. Here, we demonstrate that the use of end-modified Alt-R HDR Donor Oligos along with Alt-R HDR Enhancers, small molecules that inhibit NHEJ-mediated repair, combine to significantly increase the rate of HDR when delivered with CRISPR RNP complexes in mammalian cell lines. Altogether, this study presents a set of design considerations and reagents which can be applied to CRISPR editing experiments to maximize HDR efficiency and reduce time spent generating desired mutants. Our findings constitute an empirically defined ruleset for *S.p*. Cas9 and *S.p*. Cas9 D10A nickase which have been built into a novel bioinformatic tool for HDR donor template design. Further, we provide design recommendations for *A.s.* Cas12a nuclease, which has not yet been systematically studied in the same manner as Cas9.

## RESULTS

### Cas9 donor strand preference and gRNA selection

While some studies have suggested that there is a strand preference for the ssODN donor template, where one strand’s homology sequence consistently mediates improved HDR frequency over the other strand, results have varied and no universal strand preference has been identified.^37, 42–44^ To elucidate any universal strand preference of the HDR donor template with WT Cas9 nuclease, particularly when delivered as an ribonucleoprotein (RNP) complex, HDR efficiency was tested at 254 genomic loci in Jurkat cells and 239 genomic loci in HAP1 cells. Donor ssODNs containing 40-nt homology arms were designed to insert a six base EcoRI restriction digest recognition site (‘GAATTC’) at the Cas9 cleavage site which canonically lies three bases in the 5’ direction of the PAM, as illustrated in Figure 1A. ssODNs were delivered to Jurkat and HAP1 cells along with their respective Cas9 RNP complex by nucleofection, and the editing frequencies were assessed by next generation sequencing (NGS). Perfect HDR, defined as the precise insertion of the EcoRI sequence at the canonical cut site and otherwise maintaining the WT sequence, was quantified and comparisons were made between donor templates consisting of either the targeting strand (T), which is complementary to the CRISPR-Cas9 gRNA, or the non-targeting strand (NT), which contains the ‘NGG’ PAM sequence. In Jurkat cells there was no statistical difference (p>0.05, paired t-test) in total editing when either the T or NT strand was used. However, a significant difference in editing efficiency (p<0.0001, paired t-test) was observed in HAP1 cells where the mean editing was 80.2% when the NT strand was used and 67.8% when the T strand was used, indicating that the T strand may bind to the Cas9 RNP complex and reduce overall editing efficiency within the cellular environment, as suggested by others (Supplemental Figure 1A).^36^ As demonstrated in the top two panels of Figure 1B, the strand that leads to higher frequencies of HDR varies depending on the genomic locus and cell type being used. HAP1 cells had HDR frequencies ranging from 0 to 51.1% with a significantly higher mean HDR frequency when the NT strand was used (20.6%) than the T strand (15.2%) (p<0.0001, paired t-test), likely due to the reduced total editing when the T strand was used. In contrast, we observed significantly higher mean HDR frequencies in Jurkat cells when the T strand was used than when the NT strand was used (11.3% vs 7.5%, respectively) (p<0.0001, paired t-test). Overall, HDR efficiency in HAP1 cells was higher than in Jurkat cells, with mean HDR frequencies of 17.9% and 9.4%, respectively. In addition, the bottom two panels of Figure 1B show that although efficient total editing is required for HDR to occur, high editing does not always lead to high HDR insertion at each site tested. For example, even though 53% of the sites tested in HAP1 cells and 74% of the sites tested in Jurkat cells had >90% total editing, there is a broad range of HDR frequencies which varied from 0 to 60% among these highly edited loci for both the NT and T strands. This emphasizes the value of testing multiple guides to determine which have the highest potential HDR frequency prior to any experiment where precise genome modification by HDR is desired.

**Figure 1.**
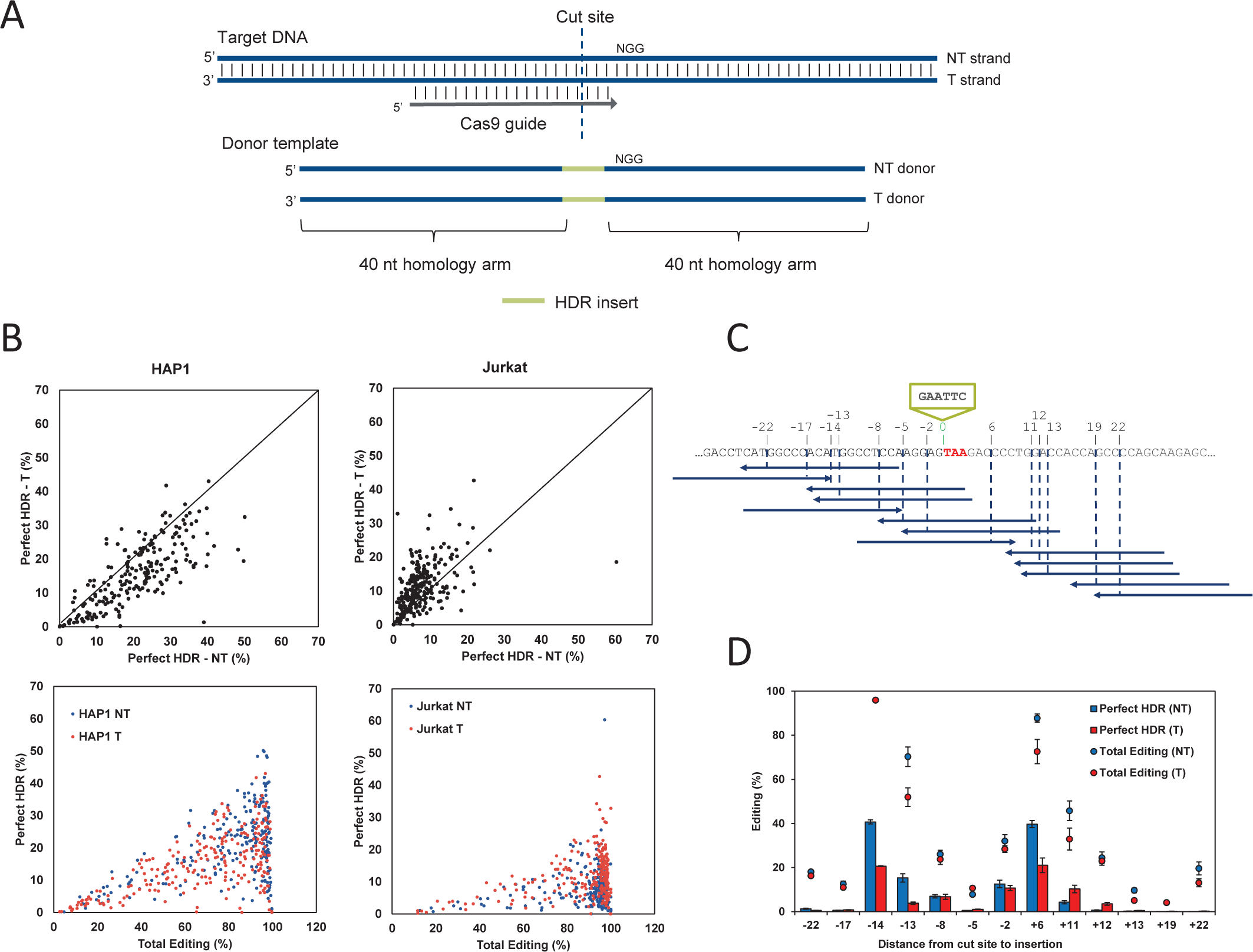
Cas9 HDR strand preference and gRNA selection. (A) Schematic representation of targeting (T) and non-targeting (NT) donor template designs. The targeting strand is complementary to the gRNA sequence, whereas the non-targeting strand contains the guide and PAM sequence (B) An EcoRI recognition site was inserted at a Cas9 cleavage site at 254 genomic loci in Jurkat and 239 genomic loci in HAP1 cells using either the T or NT strand as the donor template. RNP complexes (Alt-R S.p. Cas9 Nuclease complexed with Alt-R CRISPR-Cas9 crRNA and tracrRNA) were delivered at 4 µM along with 4 µM Alt-R Cas9 Electropo-ration Enhancer and 3 µM donor template by nucleofection. Total editing and perfect HDR was assessed via NGS. (C) Schematic of the gRNAs used to facilitate HDR insertion of an EcoRI site before the stop codon of GAPDH (TAA, red) in K562 cells using 13 guides around the desired HDR insertion location (blue, arrows indicate the 3’ end). The cleavage sites and associated distance to the desired insertion location (green) for each gRNA are indicated above the sequence shown. Both the T and NT strand were tested. (D) RNP com-plexes (Alt-R S.p. Cas9 Nuclease, Alt-R CRISPR-Cas9 crRNA and tracrRNA) for the 13 guides targeting GAPDH were delivered at 2 µM along with 2 µM Alt-R Cas9 Electroporation Enhancer and 2 µM donor template designed to insert an EcoRI site before the stop codon by nucleofection to K562 cells. HDR and total editing were assessed via NGS. Data are represented as means ± S.E.M. of three biological replicates.

To investigate the balance between guide cleavage efficiency and distance of the desired HDR mutation to the cut site, we selected 13 guides flanking the stop codon of GAPDH to determine which led to the highest HDR insertion frequency of an EcoRI site just upstream of the ‘TAA’ stop codon. These guides had cut sites that ranged from 2 to 22 bases from the desired insertion position (Figure 1C). The available guides in the nearby region included PAMs on both strands of the genomic DNA, and ssODNs for both the targeting and non-targeting strand were designed and tested in K562 and HEK293 cells for their ability to mediate HDR. As shown in Figure 1D in K562 cells, guides with low editing efficiency yielded low HDR insertion, even if the cut site was close to the desired insertion location. For example, the guide that cuts two bases from the desired insertion (-2) had 32.1% total editing of which 12.6% was HDR insertion (NT strand). Similarly, the guide that cuts five bases from the desired insertion (-5) had 10.7% total editing and only 1.1% HDR insertion. In contrast, the guides that cut 14 and 6 bases from the desired insertion (-14, +6) had 96.0% total editing of which 40.7% was HDR insertion (NT strand) and 87.8% total editing of which 39.7% was HDR insertion (NT strand), respectively. As determined by NGS, these guides had the highest total editing and HDR insertion rates, even though they were further from the desired insertion. This case study indicates that guide efficiency is a critical factor for efficient HDR, and guide selection that is as close as possible to the desired HDR mutation is a secondary consideration. This was also observed in HEK293 cells, where a guide that cuts 6 bases from the insertion (+6: 97% total editing, 34% HDR) led to higher HDR than guides 2 or 5 bases from the insertion (-5: 34.7% total editing, 12.7% HDR; -2: 62.2% total editing, 22.4% HDR). This effect was less prominent using guides further from the desired insertion site (e.g. -14) in HEK293 cells, which may be due to differences in the available repair machinery and capacity for HDR in each cell type (Supplemental Figure 1B).

We performed a similar experiment at a second genomic locus (TNPO3) in HEK293 cells to further examine factors influencing HDR (Supplemental Figure 1C). The total editing was high for nearly all guides tested in this experiment. However, for a guide with a cut site 9 bases from the desired insertion location (-9) the total editing was 92.6%, and this site yielded reduced HDR efficiency of 7.0% compared to the guide that cut one base further from the desired insertion (-10) with an increased 98.5% total editing that also gave an increased HDR insertion frequency of 24.2% with the NT strand, further supporting that guide activity can be more impactful on HDR efficiency than optimal positioning with respect to the cut site.

### Cas9 D10A mediates efficient HDR distant from nick sites

In comparison to WT Cas9, which needs only one gRNA to cut both strands of the target DNA, Cas9 D10A and H840A nickases can be used with paired guides to generate a DSB to mediate genome editing. To facilitate a DSB, the guides must target opposite strands of the genomic DNA and can be oriented with their PAM sites facing toward each other (PAM-in), or apart from each other (PAM-out) (Supplemental Figure S2A). Guides that target the same DNA strand where one PAM site would face in and the second would face out would not be generate a DSB (unless the two nickase variants, D10A and H840A, were used in combination which significantly complicates the experiment).

Consistent with other reports utilizing Cas9 nickase variants expressed from a plasmid,^17, 21^ we found a higher rate of indel formation when D10A and H840A nickases were designed in a PAM-out orientation, and the nickases must be placed with optimal spacing between the nick sites to mediate efficient editing (Supplemental Figure S2B). Here, we designed a set of paired guides against the human *HPRT1* gene with either PAM-out or PAM-in orientation and target nick sites separated by 18– 130 bp. We specifically selected gRNAs that have >40% editing efficiency (data not shown) when delivered with WT Cas9 as RNP into HEK-293 cells, to rule out the possibility that poor editing by the nickase RNP pair is caused by poor cleavage efficiency mediated by individual gRNA. The optimal distance (>40% editing as determined by T7EI) between the two nicks was 40-68 nt for Cas9 D10A, and 51-68 nt for Cas9 H840A. For the PAM-out pair with nicks 68-nt apart, the editing was 86.3% with Cas9 D10A and 77.6% with Cas9 H840A. This was reduced to 29.8% and 2.7%, respectively, when the nicks were 85-nt apart. Similarly, for Cas9 H840A, the editing was 74.7% when the nicks were spaced 51-nt apart, and this was reduced to 34.6% when the distance between the nicks was decreased to 46-nt. We have investigated distances smaller than 40-nt between the nicks in PAM-out orientation for Cas9 D10A and found that editing was poor for spacing <35-nt, likely due to steric hindrance between the two RNP molecules (data not shown).

When Cas9 D10A nickase RNP complexes targeting both strands in the PAM-out orientation nick the genomic DNA, a DSB with 5’ overhangs is generated. Because both strands are targeted by one of the two gRNAs, there is no canonical ‘targeting’ and ‘non-targeting’ strand in nickase experiments; thus, they are referred to as top and bottom strands. The paired-guide double nicking strategy doesn’t generate a blunt-ended cut like WT Cas9, so we further explored the possibility of using Cas9 D10A to insert exogenous sequences between flanking nick sites at a location that would be otherwise considered sub-optimal for WT Cas9 designs using WT Cas9 and either gRNA on its own. We designed ssODN donor templates for *HPRT1* in which a 6-nt EcoRI restriction enzyme recognition site was introduced at different locations along a donor template (Figure 2A). HDR events mediated by Cas9 D10A in HEK293 cells, as measured by the percentage of EcoRI digestion, ranged from 13-25% across the 51-nt region (Figure 2B, top left panel). In contrast, WT Cas9-mediated HDR decreased dramatically as the intended insertion site moved away from the cleavage site (Figure 2B, top middle and right panels), consistent with our earlier findings. At the position centered between the two cleavage sites (25-nt from left and 26-nt from the right), Cas9 D10A was able to induce a higher HDR insertion frequency than WT Cas9 with either of the individual gRNAs. As a comparison, neither WT Cas9 nor Cas9 H840A with the same gRNA pair demonstrated HDR insertion frequency as high as Cas9 D10A at all positions tested (Supplementary Figure S2C). In addition, at these sites WT Cas9 demonstrated a strong preference for the NT strand donor template (bottom strand for WT with left gRNA, and top strand for WT with right gRNA), especially at positions distant from the cut site, while Cas9 D10A did not show a strand preference. We performed the same experiment in K562 cells, which demonstrated robust HDR overall with WT Cas9. As expected, despite both Cas9 D10A and WT Cas9 showing a higher frequency of HDR in this cell line, the ability of Cas9 D10A to mediate higher HDR when moving away from cleavage sites was retained (Figure 2B, bottom panels).

**Figure 2.**
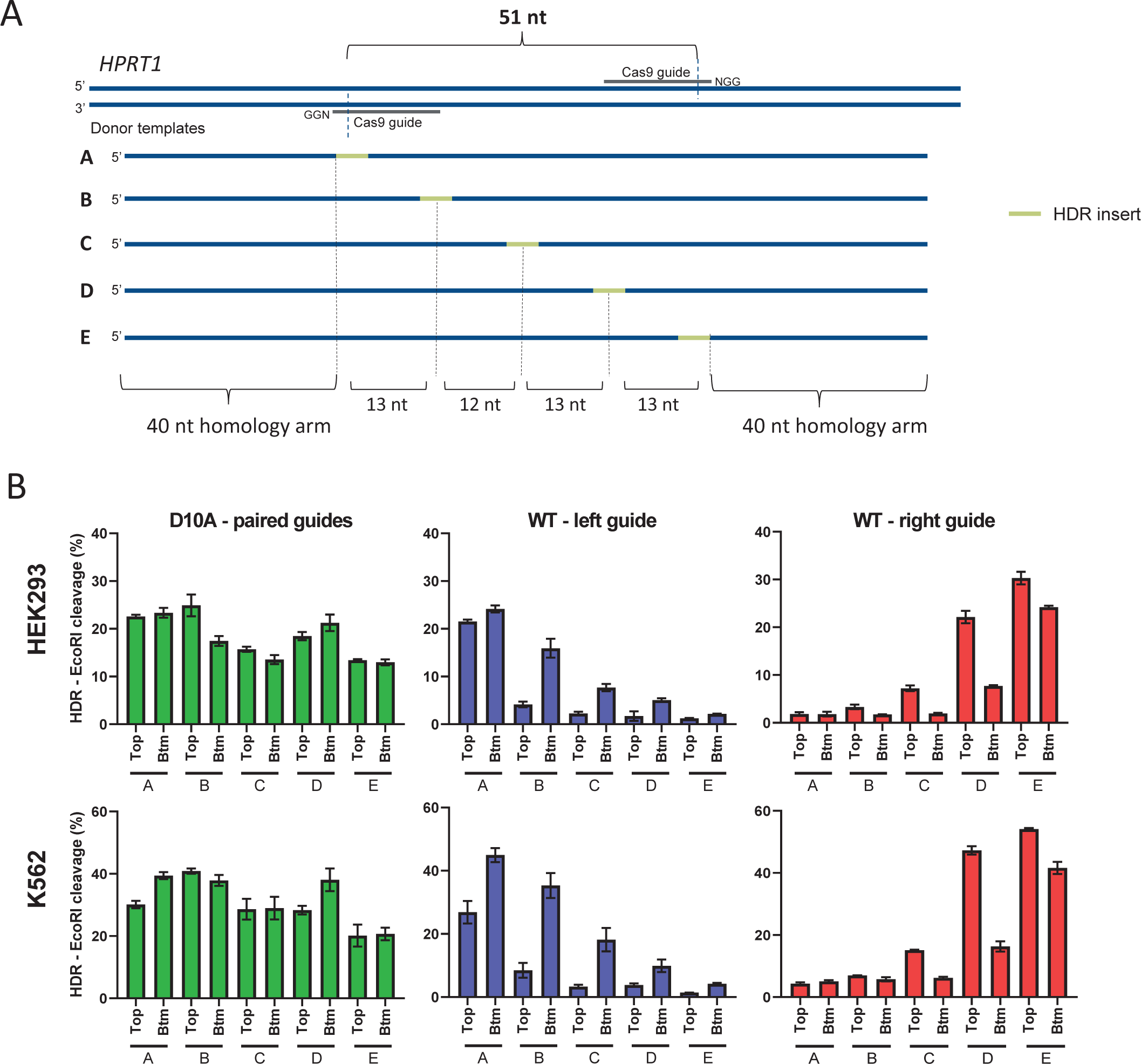
Cas9 D10A mediates efficient HDR distant from nick sites. (A) Ten HDR donor templates were designed with an EcoRI sequence positioned at varying distances (0-nt, 13-nt, 25-nt, 38-nt and 51-nt) from the left cleavage site of a paired-guide nickase design with a PAM-out orientation in HPRT1. ssODNs corresponding to the top and bottom (Btm) strand for each sequence were tested. Letters A-E indicate the position of the EcoRI insertion, the top strand ssODN is shown. (B) Cas9 D10A with gRNA pairs (left panel), or Cas9 WT with each of the individual gRNAs (middle and right panel) RNP complexes (Alt-R S.p. Cas9 D10A nickase or Alt-R S.p. Cas9 Nuclease complexed with Alt-R CRISPR-Cas9 crRNA and tracrRNA) were delivered at 4 µM (2 µM each RNP for nickase paired guides) along with 4 µM Alt-R Cas9 Electroporation Enhancer and 2 µM donor template by nucleofection to HEK293 cells (top) or K562 cells (bottom). HDR efficiency was evaluated by EcoRI cleavage of targeted amplicons. Data are represented as means ± S.E.M of three biological replicates for D10A and two biological replicates for WT.

In order to verify that the above observations are not site specific, we conducted a similar experiment at a different locus (*AAVS1*, PAM-out design with 46-nt spacing). In addition to 5 insert positions at or between the two cleavage sites, we also included two positions 12-nt upstream or downstream to the left or right cleavage sites, respectively (Supplementary Figure S2D). Consistent with the results described above, Cas9 D10A outperformed WT Cas9 at the position centered between the cleavage sites (position “D”) in HEK293 cells (Supplementary Figure S2E). Insertions outside of the nick sites were not as efficient as ones placed between the nick sites. The observation that Cas9 D10A has no identifiable strand preference for the HDR donor template also held true at this site. For HDR experiments with Cas9 D10A, the highest editing efficiency occurred when paired gRNAs were in the PAM-out orientation with the nick sites spaced 40-68 nt apart. Overall, we observed that the desired mutation is best achieved when placed between the two nicks, and we did not observe a consistent strand preference. The use of paired gRNAs with Cas9 D10A nickase allows for HDR insertions at locations not accessible by WT Cas9 nuclease due to design limitations and may be advantageous over WT Cas9 in situations where there is a lack of efficient Cas9 guides near the intended HDR mutation.

### Optimizing placement and number of blocking mutations with Cas9

Previous studies have demonstrated that incorporating blocking mutations within the donor oligo to prevent re-cleavage by Cas9 nuclease after a desired HDR event improves rates of HDR.^35, 45^ However, these studies have been limited in the number of constructs tested and have not examined if there is a preference for transversions (e.g. G-to-C purine to pyrimidine conversion), or transitions (e.g. G-to-A purine to purine conversion) in the blocking mutations used. We aimed to further investigate this to define a ruleset for the placement and number of blocking mutation(s) required to maximize HDR efficiency. First, we designed an experiment to determine the effect of a single blocking mutation within the PAM or the seed region of Cas9, which is defined as the PAM-proximal 10-12 bases on the 3’ end of the guide.^7^ Mismatches within the seed region and PAM are known to significantly reduce Cas9 binding and cleavage efficiency, so would be expected to confer the highest reduction in re-cleavage by Cas9.^8, 50^ Two genomic loci were selected and HDR ssODN donor templates were designed to generate a single base change 3’ of the PAM to serve as the desired HDR mutation. This HDR mutation would not impact Cas9 re-cleavage, as it falls outside of the protospacer/PAM sequence. In addition to the desired HDR mutation, a single blocking mutation in the seed region of the guide or PAM was included, where each position tested was changed to every possible alternate base in a unique donor template to determine if any of the four DNA bases are preferred when utilizing blocking mutations (Figure 3A). HDR ssODN donor templates were delivered along with their respective RNP complexes targeting two different genomic loci into HEK293 and K562 cells, and the rate of perfect HDR including both the desired HDR mutation and blocking mutation (where applicable) was assessed by NGS (Figure 3B). As expected, the frequency of the desired HDR mutation (‘ctrl’) was low, at <2% for all four conditions tested. Adding a blocking mutation in the second or third base of the ‘NGG’ PAM resulted in the greatest increase, with HDR levels reaching 8.0-17.8%. Blocking mutations placed around the Cas9 cleavage site and near the 3’ end of the guide were also highly effective, with the impact reduced as the position of the blocking mutation moved PAM-distal. No base was universally preferred over others in these experiments. Next, we aimed to determine if a single blocking mutation was sufficient to prevent re-cleavage of the genomic DNA and maximize the number of HDR events, or if a combination of multiple blocking mutations would lead to higher HDR frequency. Donor templates were designed with a single blocking mutation in the PAM, two blocking mutations in the PAM, and/or blocking mutations at two locations within the seed region. Various combinations of these mutations were delivered as ssODNs along with corresponding RNP complexes targeting four loci in Jurkat cells to assess their ability to mediate a single base change 3’ of the PAM via HDR. An example sequence showing the placement of blocking mutation(s) in the donor templates tested is provided in Supplemental Figure 3A. As demonstrated by Supplemental Figure 3B, donor templates containing two blocking mutations led to more robust improvement in HDR efficiency than donor templates containing a single blocking mutation, and this effect was greatest when the blocking mutations were within the PAM or nearer to the 3’ end of the guide. Incorporating three or four blocking mutations did not further enhance HDR efficiency over the best combination of 2 blocking mutations (2 PAM or 1 PAM + 1 seed A).

**Figure 3.**
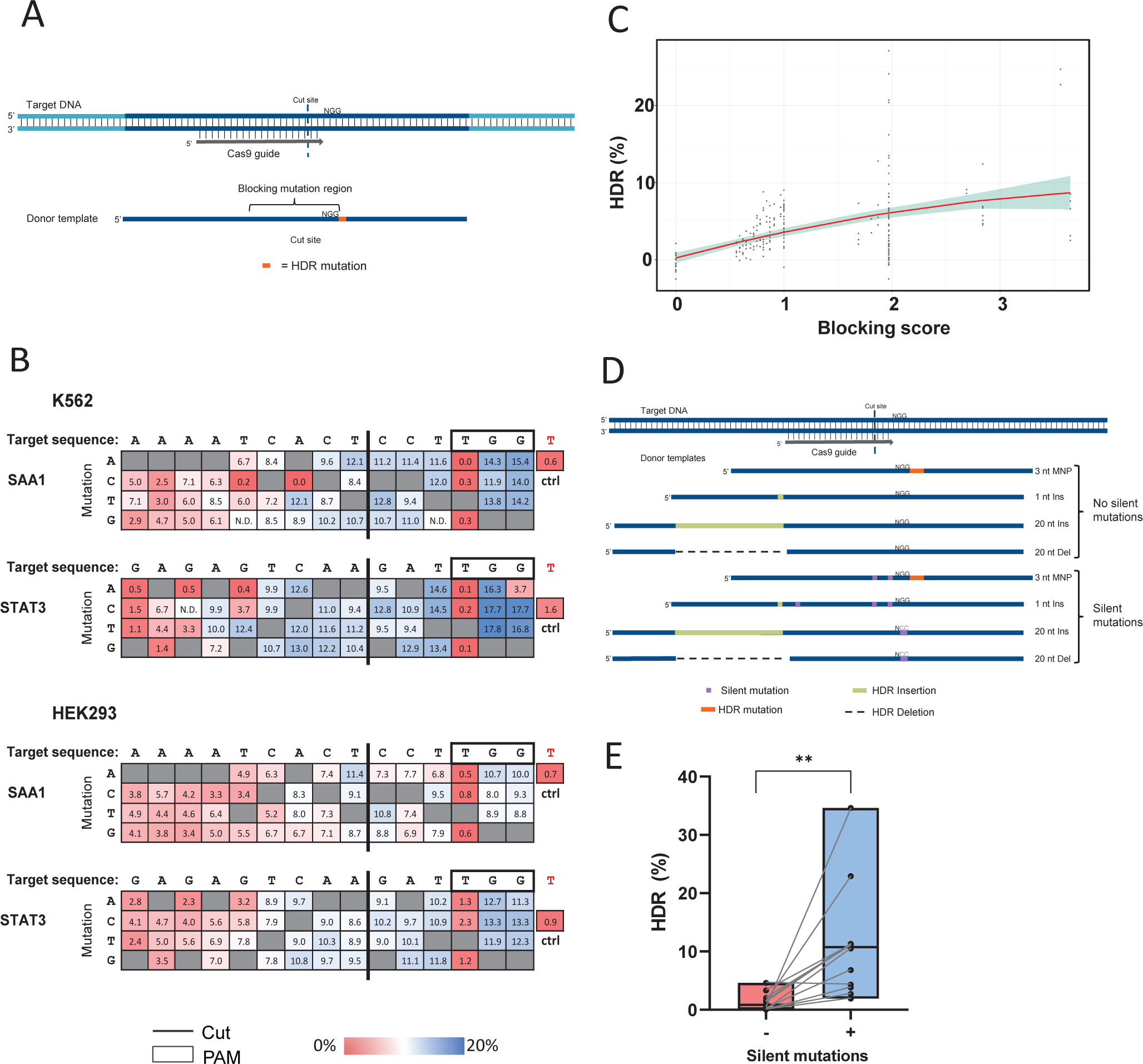
Optimizing placement and number of blocking mutations with Cas9. (A) Schematic representation of a desired HDR generated single base change (orange) 3’ of the PAM. In addition to the desired HDR mutation, a single blocking mutation in the seed region of the guide or PAM to prevent Cas9 re-cleavage was included in donor templates. Each position in the region indicated was changed to every possible alternate base in a unique donor template that also contained the desired HDR mutation. (B) HDR donors for two genomic loci were tested in HEK293 and K562 cells. In each case the donor contained an HDR mutation 3’ of the PAM, with or without a blocking mutation within the region indicated. HDR donors were delivered at 4 µM along with RNP complexes (Alt-R S.p. Cas9 Nuclease complexed with Alt-R CRISPR-Cas9 crRNA and tracrRNA) at 4 µM and with 4 µM Alt-R Cas9 Electroporation Enhancer by nucleofection. Each box represents the rate of perfect HDR including both the desired HDR mutation and blocking mutation (where applicable) as assessed by NGS. Blue indicates a higher HDR frequency, and red indicates a lower HDR frequency. (C) Blocking scores were calculated for 427 samples with known HDR frequencies and used to build a linear model (model = red line, standard error = blue highlight) to determine the optimum HDR efficiency. (D) Schematic representation of four unique HDR mutations that were designed using the Alt-R HDR Design tool either with or without the addition of silent mutations. (E) Four HDR mutations designed using the novel Alt-R HDR Design Tool with (+) or without (-) silent mutations were tested in HEK293, Hela, and Jurkat cells. RNP complexes (Alt-R S.p. Cas9 Nuclease, Alt-R CRISPR-Cas9 crRNA and tracrRNA) were delivered at 4 µM along with 4 µM Alt-R Electroporation Enhancer and 4 µM donor template in HEK293 and Jurkat cells by nucleofection. RNP complexes (Alt-R S.p. Cas9 Nuclease complexed with Alt-R CRISPR-Cas9 sgRNA) were delivered at 2 µM along with 2 µM Alt-R Cas9 Electroporation Enhancer and 2 µM donor template in Hela cells by nucleofection. Perfect HDR rates were determined by NGS.

We next wanted to investigate this effect when a larger HDR mutation is inserted, such as an EcoRI restriction site, as well as examine the impact of blocking mutations when the HDR insertion was placed at various positions relative to the Cas9 cleavage site. To determine if blocking mutations are beneficial with a 6-nt insertion, we selected four gRNAs and designed donor templates to insert an EcoRI restriction digest recognition site at the Cas9 cleavage site. Donor templates included no blocking mutation (no PAM mutation) or a ’GG’ to ‘CC’ blocking mutation within the PAM sequence (PAM mutation). In addition, donor templates to insert the EcoRI sequence at varying locations relative to the Cas9 cleavage were designed; as a result, these donor templates would facilitate the 6-nt insertion as close as 3-nt from the Cas9 cleavage site and extending as far as 45-nt in both the 5’ and 3’ direction. Designs of donor templates again contained no blocking mutation or a ‘GG’ to ‘CC’ PAM mutation (Supplemental Figure 3C). All donor templates consisted of the NT-strand and maintained homology arms of 40-nt from both the EcoRI insertion location and the Cas9 cut site. The set of 24-32 ssODNs for the four targets were delivered along with their respective Cas9 RNP complex to Jurkat cells by nucleofection (N = 120). Additionally, donor templates and respective Cas9 RNP complexes for two of the targets were also delivered to HEK293 cells (N = 48). An EcoRI cleavage assay was used to determine the HDR frequencies, and results are shown in Supplemental Figure 3D. Mutating the PAM from an ‘NGG’ to ‘NCC’ increased HDR at locations further from the cut site in the 3’ direction, downstream of the PAM and outside of the protospacer sequence, where the HDR insertion would not prevent Cas9 re-cleavage. When the EcoRI insertion was within the guide seed region or PAM, incorporating additional PAM mutations negatively impacted the HDR efficiency. This suggests that there is a limit to the number of additional mutations that should be added to prevent Cas9 re-cleavage, and if too many mutations are present the HDR efficiency can be negatively affected.

This data represents a subset of HDR donor designs that we tested to fully elucidate a ruleset for the placement and number of blocking mutations required for various HDR mutation types. Using HDR efficiency results from Figure 3B, we generated relative HDR efficiencies (i.e., HDR efficiency with varying blocking mutations divided by HDR efficiency without blocking mutations) and a position specific scoring matrix (PSSM). The PSSM represents the HDR improvement introduced by mutating the HDR donor template at each position along the length of the Cas9 spacer sequence. Using the linear combination PSSM values representing blocking mutations in each HDR donor template, we calculated a blocking score for each of 374 donor template designs associated with 9 guides and delivered into 3 cell lines (HEK293, Jurkat, and Hepa1-6). We generated a model representing a non-linear correlation between blocking scores and HDR efficiency (Figure 3C). A score of 1.97 approximately corresponds to mutating both G nucleotides in the Cas9 PAM. The model predicts blocking mutations with scores <1.97 will have a positive impact on HDR rates; while blocking mutations with scores greater than 1.97 have a less certain, and perhaps detrimental, impact. We embedded the PSSM, blocking score model, a guide-to-target mutation model, and other heuristics in the Alt-R HDR Design Tool. The combination of models allows the tool to recommend high quality paired HDR donor templates and guides. In addition, the Alt-R HDR Design Tool uses blocking scores to select block mutations that do not change the protein coding sequence (when transcript information is provided). We tested the Alt-R HDR Design Tool’s donor template recommendations using four unique target HDR mutations with or without the addition of silent blocking mutations (Figure 3D), and we delivered the donor templates to HEK293, Hela and Jurkat cells. In every case except one in Jurkat cells, where the HDR rate was unchanged, the donor template designed with the addition of silent mutations yielded higher HDR events than donor template designs without blocking silent mutations (Figure 3E).

### HDR mutation location determines donor strand preference

Achieving efficient HDR at greater distances from the cut site using WT Cas9 to broaden the capabilities of CRISPR genome editing is desirable. We aimed to investigate design considerations for HDR mutations that fall outside of the optimal editing window to determine if there is a donor strand preference. Additionally, we tested if additional mutations along the ssODN repair track, which is defined as the portion of the donor template between the Cas9 cut site and mutation location, were beneficial. We designed ssODN donor templates to create an EcoRI insert 25-nt from the Cas9 cleavage site either on the PAM-containing side of the Cas9 cut (PAM-proximal) or on the non-PAM side of the Cas9 cut (PAM-distal) for three genomic loci. Donor templates were designed to have 1) no mutation, 2) an ‘NGG’ to ‘NCC’ PAM mutation to prevent re-cleavage after HDR, or 3) mutations placed along the repair track every 5^th^ nt, either alone or in combination with the PAM mutations (Figure 4A). Both the T and NT strands were tested to determine which donor templates facilitated the highest HDR incorporation of an insert outside of the previously established optimal placement. ssODN donor templates were delivered along with their respective Cas9 RNP complexes to HeLa cells by nucleofection, and the frequency of perfect HDR containing both the desired HDR mutation and any additional mutations was determined by NGS. The mean HDR rate for each ssODN design across three biological replicates for each of the three genomic loci tested is shown in Figure 4B (for each ssODN design n = 9). For PAM-distal insertions, the NT strand had an average HDR of 12.7% with repair track mutations compared to 1.6% when the T strand was used. In contrast, for PAM-proximal insertions, the T strand containing repair track and PAM mutations gave higher HDR than the NT strand with the same mutations (8.6% vs 0.8%, respectively). For PAM-proximal insertions, the repair track mutations marginally improved the HDR efficiency above incorporating a PAM mutation alone, increasing the HDR from 7.5% to 8.6%. However, for PAM-distal insertions, the repair track mutations significantly improved the frequency of HDR 3.4-fold over having only a PAM mutation (p <0.01, paired t-test).

**Figure 4.**
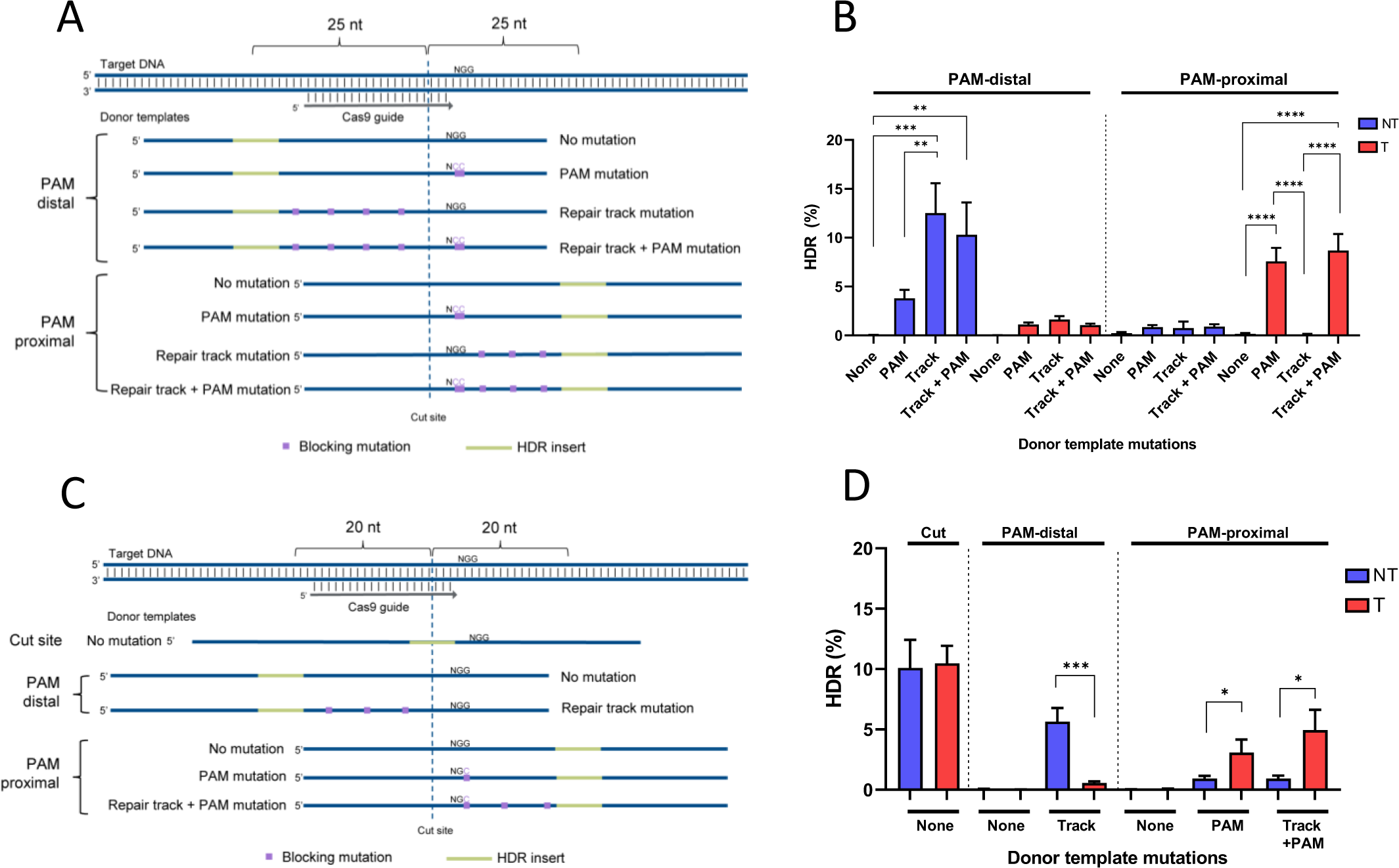
HDR mutation location determines donor strand preference. (A) Schematic representation of donor templates used to generate PAM-proximal and PAM-distal insertions 25 bases from a Cas9 cut site with no further mutations (None), PAM mutations (PAM), or mutations in the repair track with or without an additional PAM mutation. The NT strand ssODNs are shown. (B) Donor templates creating an EcoRI insertion 25 bases from the cut site at three genomic loci were delivered to Hela cells as the T or NT strand. Donor templates contained no further mutation (None), PAM mutation (PAM), or mutations in the repair track (Track). RNP complexes (Alt-R *S.p.* Cas9 Nuclease complexed with Alt-R CRISPR-Cas9 sgRNA) were delivered at 2 µM along with 2 µM Alt-R Cas9 Electroporation Enhancer and 0.5 µM donor template by nucleofection. Perfect HDR rates were determined by NGS. Data are represented as means ± S.E.M of the three sites tested. (C) Schematic representation of donor templates used to generate PAM-proximal and PAM-distal insertions 20 bases from a Cas9 cut site with no further mutations (None), PAM mutations (PAM), or mutations in the repair track with or without additional PAM mutation. The NT strand ssODNs are shown. (D) Donor templates creating an EcoRI insertion at the cut site or 20 bases PAM-proximal or PAM-distal to the Cas9 cut site for 12 genomic loci were tested in Jurkat cells as the T or NT strand. RNP complexes (Alt-R *S.p*. Cas9 Nuclease complexed with Alt-R CRISPR-Cas9 sgRNA) were delivered at 4 µM along with 4 µM Alt-R Cas9 Electroporation Enhancer and 3 µM donor template by nucleofection. Perfect HDR rates were determined by NGS. Data are represented as means ± S.E.M of the twelve sites tested.

To further investigate strand preference when HDR mutations are placed at suboptimal distances (>15-nt) away from the Cas9 cleavage site, 12 loci from the set of 254 targets presented in Figure 1B were selected as a subset of gRNAs to carry out this experiment. These gRNAs were selected as sites for HDR because they demonstrated one of three characteristics: no strand preference, an obvious strand preference for the T strand, or an obvious strand preference for the NT strand in either Jurkat or HAP1 cells. Donor templates were designed following the same principles as the prior experiment, placing an EcoRI insertion at the Cas9 cleavage site or 20 bases PAM-proximal or PAM-distal. Given the results shown in Figure 4B that identify PAM-distal insertions as mediating sub-optimal insertion frequencies, a single PAM mutation alone was only tested for PAM-proximal insertions. In addition, repair track mutations were incorporated every 3-7 nt between the Cas9 cleavage site and the desired HDR mutation (Figure 4C). These ssODN donor templates were delivered to Jurkat and HeLa cells along with their respective Cas9 RNP complexes by nucleofection, and the frequency of perfect HDR was determined by NGS with the mean HDR rate for each ssODN across the 12 genomic loci shown in Figure 4D. Across all 12 sites tested, the NT strand gave higher HDR than the T strand for PAM-distal insertions, and the T strand gave higher HDR than the NT strand for PAM-proximal insertions (Figure 4D, Supplemental Figure 4A). Similar to the previous experiment, repair track mutations in combination with a single PAM mutation for PAM-proximal insertions had a modest improvement in HDR rates over the single PAM mutation alone, increasing from 3.0% to 4.9% in Jurkat cells and 5.4% to 8.7% in Hela cells. The level of HDR improvement for the various mutation strategies had site-to-site variability (Supplemental Figure 4B). However, the strand preference was universal to all sites tested, indicating that for PAM-proximal insertions the T strand should be used as the donor template, and for PAM-distal insertions, the NT strand should be used for the highest rate of HDR.

### Optimized design rules for HDR with Cas12a

Cas12a is a type II CRISPR-Cas nuclease with several distinct differences to Cas9. Cas12a generates a DSB with 5’ overhangs, requires a ‘TTTV’ PAM, and enables editing in AT-rich genomes.^4^ We designed experiments to characterize HDR design rules for Cas12a in a manner similar to what was done with Cas9. First, the optimal placement of an insertion was determined by designing donor templates for five sites in the *HPRT1* gene. These donor templates placed an EcoRI restriction digest recognition site at varying positions relative to the PAM and guide sequence (Figure 5A), ranging from 9 bases away in the 5’ direction from the first base of the guide to 45 bases 3’ of the first base of the guide. The optimal HDR activity for this insert is not centered around the two Cas12a cleavage sites, canonically positioned 18 and 23 bases from the PAM, as was the case for Cas9. There is a strong preference for insertions between positions 12-16 of the guide (Figure 5B).

**Figure 5.**
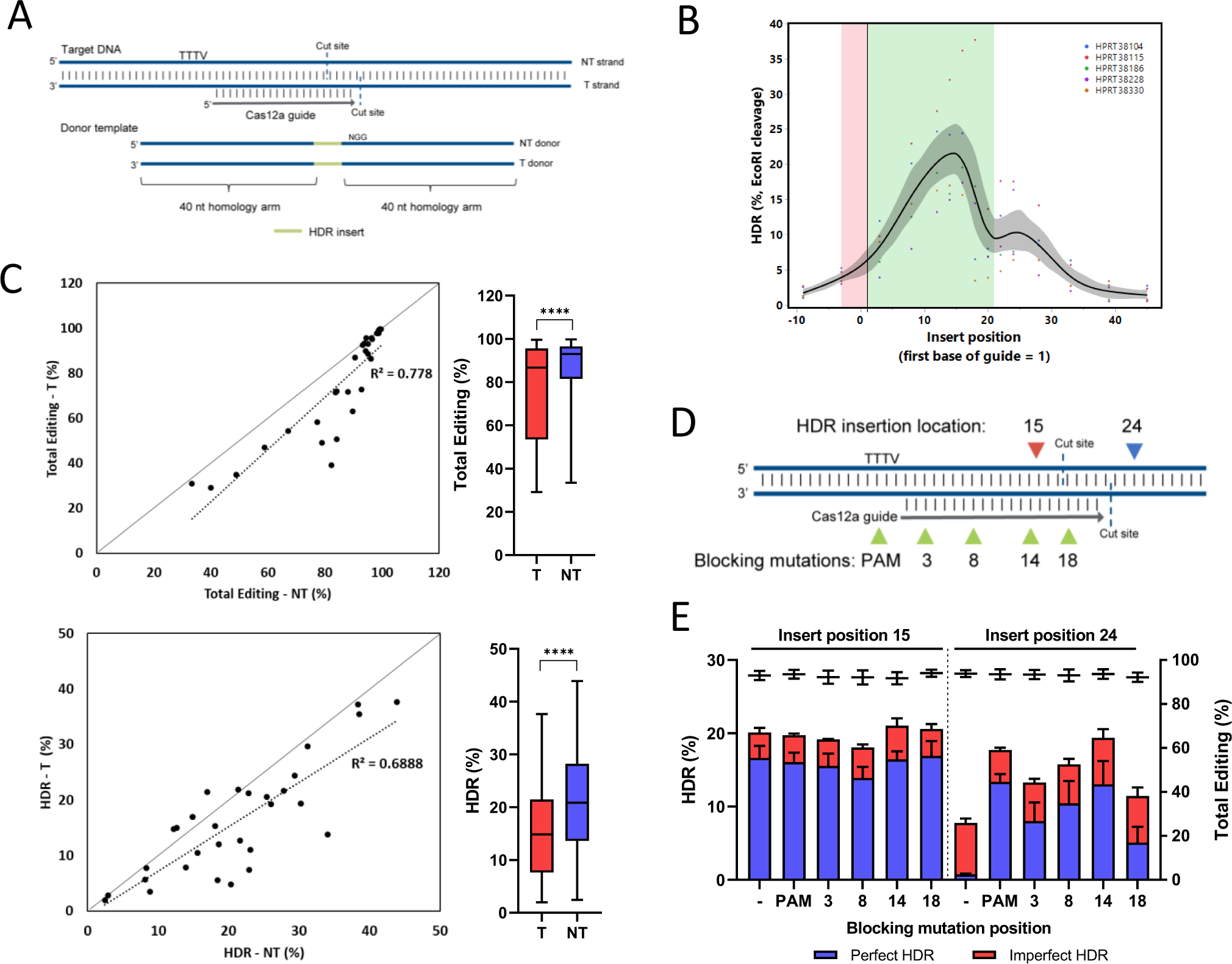
Cas12a HDR gRNA selection and strand preference. (A) Schematic representation of targeting (T) and non-targeting (NT) donor template designs. The T strand is complementary to the gRNA sequence, whereas the NT strand contains the guide and PAM sequence. (B) HDR donors were designed with an EcoRI insert sequence positioned at varying distances from the from the first base of the Cas12a guide RNA ranging from 10 bases in the 5’ direction to 45 bases in the 3’ direction for five genomic loci and delivered to HEK293 cells. RNP complexes (Alt-R *A.s.* Cas12a nuclease complexed with Alt-R CRISPR-Cas12a crRNA) were delivered at 5 µM along with 3 µM Alt-R Cpf1 Electroporation Enhancer and 3 µM donor template by nucleofection. HDR rates were assessed via EcoRI cleavage of targeted amplicons. The 21-bases where the gRNA targets is highlighted in green. The 4 base ‘TTTV’ PAM is highlighted in red. The gray shading indicates the confidence of fit. (C) An EcoRI restriction digest recognition site was inserted at position 16 of the gRNA sequence in 15 genomic loci in Jurkat and HAP1 cells using either the T or NT strand as the donor template and the combined results graphed together. RNP complexes (Alt-R *A.s.* Cas12a *Ultra* nuclease complexed with Alt-R CRISPR-Cas12a crRNA) were delivered at 1 µM along with 3 µM Alt-R Cpf1 Electroporation Enhancer and 3 µM donor template by nucleofection. Total editing and perfect HDR was assessed via NGS. (D) Donors for two genomic loci were designed to insert an EcoRI site within the Cas12a guide sequence (position 15 of the guide) or outside of the guide sequence (24 bases from the start of the guide). ssODNs for these two insert locations were designed with blocking mutations in the PAM or guide sequence. The positions where blocking mutations were incorporated are indicated. (E) Donor templates for two genomic loci in *HPRT1* were tested in Jurkat and Hela cells. RNP complexes (Alt-R *A.s.* Cas12a *Ultra* complexed with Alt-R CRISPR-Cas12a crRNA) were delivered at 2 µM along with 2 µM Alt-R Cpf1 Electroporation Enhancer and 3 µM donor template by nucleofection. HDR rates were assessed via NGS. Perfect HDR (blue), imperfect HDR (red) and total editing, which includes NHEJ events (black) are shown. Data are represented as means ± S.E.M.

Interestingly, there is an increase in EcoRI insertion around position 24 even though this position falls outside of the protospacer region. We hypothesized this to be a result of imperfect HDR where an EcoRI site is inserted via HDR, followed by Cas12a re-cleavage which then allows the insertion of other indels from NHEJ repair. To investigate this possibility, we performed NGS analysis of one of the five sites from Figure 5B to examine the frequency of perfect HDR insertion relative to imperfect HDR insertion. At position 24, while the amount of EcoRI insertion was 6.4% by EcoRI cleavage (Supplemental Figure 5A), the amount of perfect HDR when measured by NGS is <1% and the imperfect HDR, which includes HDR insertion of an EcoRI site plus subsequent indels from NHEJ due to Cas12a re-cleavage, was 5.9% (Supplemental Figure 5B). Thus, we confirmed by NGS that the optimal position for Cas12a-mediated HDR is between positions 12-16 of the guide and moving an insertion outside of the protospacer can give the desired insertion, but also allows for additional undesired editing.

To investigate if Cas12a demonstrates a universal strand preference when an EcoRI insertion was optimally placed, a set of 15 Cas12a guide RNAs was selected and donor templates were designed to insert an EcoRI restriction digest recognition site 16 bases 3’ of the PAM. Both the T and NT strand ssODN donor templates were delivered with their respective RNP complexes to Jurkat and HAP1 cells by nucleofection, and NGS was used to measure the frequency of total editing and perfect HDR. The combined results from the fifteen sites comparing T and NT strand donors in two cell lines is shown in Figure 5C. Although there are differences in total editing across the 15 sites tested (varying from 30% to >95% total editing which indicates inherent, guide or locus-dependent editing outcomes), universally the total editing was lower when the T strand was used. This is shown in figure 5C, top panel by the data points generally clustering below the line through the origin or showing increased total editing when delivered with the NT strand. The reference line through the origin is included as a benchmark in both panels for 5C to indicate the point at which T strand total editing or HDR is equivalent to NT strand total editing or HDR, respectively. As a result of the discrepancy observed in favor of the NT strand mediating increased total editing (top panel of 5C), the frequency of HDR was also lower when the T strand was used as the donor template than when the NT strand was used. These results demonstrate a statistically significant preference for the use of the NT strand as the donor template to achieve optimal results in HDR experiments using Cas12a nuclease.

The results from Figure 5B and follow-up in Supplemental Figure 5B suggest that blocking mutations could also be beneficial in Cas12a-mediated HDR. We designed experiments to investigate whether HDR could be improved at a position outside of the guide targeting sequence by incorporating blocking mutations within the ssODN donor template. Donor templates with an EcoRI insertion optimally placed at position 15 of the guide, or sub-optimally at position 24 from the first base of the guide (outside of the guide targeting region) were designed to include no blocking mutation, a blocking mutation of the PAM sequence (TTTV to TVTV), or a blocking mutation within the guide targeting sequence at various positions (Figure 5D). These were tested as NT strand donor templates at two genomic loci within the *HPRT1* gene and in two cell lines, Jurkat and Hela. When the EcoRI cleavage site was inserted within the guide sequence there was no benefit to including blocking mutations to prevent further re-cleavage, likely because the EcoRI site disrupts subsequent cleavage events (Fig 5E, left panel). However, when the EcoRI insertion was outside of the PAM/guide targeting region, blocking mutations increased the rate of HDR from 0.7%, to 13.3% with a mutation in the PAM and 13.0% with a mutation at position 14 of the guide (Figure 5E, right panel). These results show that, similar to Cas9, blocking mutations are beneficial with Cas12a and can be used to broaden the available window for efficient HDR insertions.

### Alt-R modified HDR Donor oligos and Alt-R HDR Enhancer reagents further improve HDR

It has been previously reported that the addition of phosphorothioate (PS) modifications improve HDR efficiency.^39, 46^ We investigated over 20 different stabilizing modifications (data not shown) and have developed Alt-R HDR Donor Oligos which include 2 PS linkages at the ultimate and penultimate backbone linkage and an end-blocking modification at both the 5’ and 3’ end to provide increased stability. We designed 7 donor templates to insert a 6-nt EcoRI site with 30 to 40-nt homology arms at unique genomic loci, and 1 donor template designed to insert a 42-nt sequence with 60-nt homology arms. These contained either no modification (unmodified), two PS linkages at each end of the donor template (PS modified) or Alt-R modified donor templates. They were delivered to HeLa cells by nucleofection along with corresponding RNP complexes consisting of sgRNAs complexed with Alt-R *S.p.* HiFi Cas9 nuclease. PS modified donor templates confer improved HDR over unmodified donor templates, with an average of 3.1-fold improvement (Figure 6A). Alt-R modifications provided further increase in HDR over both unmodified and PS modified donor templates by an average of 5.2-fold and 1.7-fold, respectively.

**Figure 6.**
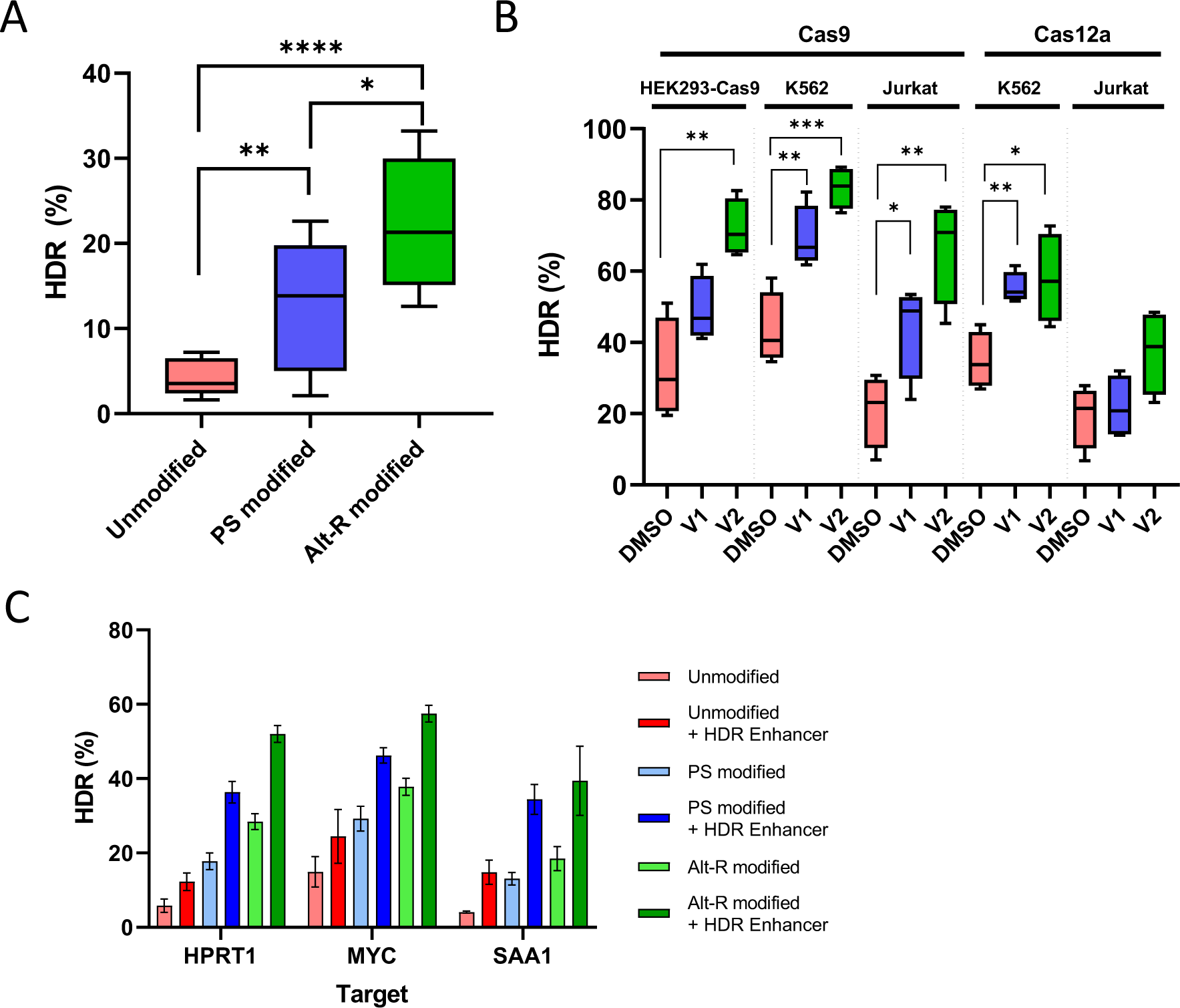
Alt-R modified HDR Donor oligos and Alt-R HDR Enhancer reagents further improve HDR. (A) Hela cells were transfected with 2 µM Cas9 RNP complexes (Alt-R *S.p.* HiFi Cas9 Nuclease complexed with Alt-R CRISPR-Cas9 sgRNA) targeting 8 genomic loci along with 0.5 µM HDR donor template and 2 µM Alt-R Cas9 Electroporation Enhancer by nucleofection. Donor templates contained no modifications (Unmodified), 2 phosphorothioate linkages between the first and last three bases of the template (PS modified), or the Alt-R HDR modification (Alt-R modified). HDR efficiency was measured by NGS. (B) HDR donor templates for four genomic loci for Cas9 and four genomic loci for Cas12a were designed to insert an EcoRI site at the cut site (Cas9) or at the 16^th^ base of the guide (Cas12a). To test lipofection delivery, gRNA complexes were delivered at 10 nM gRNA (Alt-R Cas9 crRNA and tracrRNA) with 3 nM donor template into HEK293-Cas9 cells. For Cas9 sites, K562 and Jurkat cells were transfected with 2 µM RNP (Alt-R *S.p*. Cas9 Nuclease complexed with Alt-R CRISPR-Cas9 crRNA and tracrRNA), 3 µM Alt-R Cas9 Electroporation Enhancer, and 3 µM donor template. For Cas12a sites, K562 and Jurkat cells were transfected with 2 µM RNP (Alt-R *A.s.* Cas12a *Ultra* complexed with Alt-R CRISPR-Cas12a crRNA), 3 µM Alt-R Cpf1 Electroporation Enhancer, and 3 µM donor template. Immediately after transfection, cells were plated in media containing a DMSO control, 30 µM Alt-R HDR Enhancer V1, or 1 µM Alt-R HDR Enhancer V2 and media was changed after 24 hours. HDR efficiency was measured by NGS. (C) Hela cells were transfected with 2 µM Cas9 RNP complexes (Alt-R *S.p*. HiFi Cas9 Nuclease complexed with Alt-R CRISPR-Cas9 crRNA and tracrRNA) targeting 3 genomic loci along with 0.5 µM HDR donor template and 2 µM Alt-R Cas9 Electroporation Enhancer by nucleofection. Donor templates were unmodified, PS modified, or Alt-R modified. Immediately after electroporation, cells were plated in media with or without 30 µM Alt-R HDR Enhancer (V1) and media was changed after 24 hours. HDR efficiency was measured by NGS. Data are represented as means ± S.E.M of three biological replicates.

Another strategy to increase HDR is to add chemical compounds that inhibit NHEJ repair and promote HDR repair.^47–49^ We tested two commercially available compounds, Alt-R HDR Enhancer V1 and Alt-R HDR Enhancer V2, for their ability to promote HDR. Four genomic loci for Cas9 and four genomic loci for Cas12a were selected and Alt-R modified donor templates containing 40-nt homology arms were designed to insert an EcoRI cleavage site at the optimal position (at the Cas9 cleavage site or at position 16 of the Cas12a guide). After standard delivery of ssODNs with gRNAs via lipofection (HEK293-Cas9 cells, which stably express Cas9 nuclease) or ssODNs with RNP complexes by nucleofection (K562, Jurkat) cells were plated in media containing HDR Enhancer V1, HDR Enhancer V2, DMSO, or untreated (data not shown), with a media change after 24 hours to standard media. With Cas9, HDR Enhancer V1 increased HDR frequencies by 1.5-, 1.6-, and 2.1-fold in HEK293-Cas9, K562, and Jurkat cells, respectively. With Cas12a delivery, HDR Enhancer V1 increased HDR frequencies 1.6, and 1.1-fold in K562 and Jurkat cells, respectively. With Cas9, HDR Enhancer V2 increased HDR frequencies 2.2-, 1.9-, and 3.2-fold in HEK293-Cas9, K562, and Jurkat cells, respectively. With Cas12a delivery, HDR Enhancer V2 increased HDR frequencies 1.7- and 1.9- fold in K562 and Jurkat cells, respectively (Figure 6B).

Finally, we investigated if the use Alt-R HDR Enhancer V1 in combination with modified donor templates improved HDR further than if just one of these reagents was used. In this experiment Alt-R HDR Enhancer V1 was used, although we have observed similar results with Alt-R HDR Enhancer V2 (data not shown). We found that the maximal HDR efficiency was achieved when Alt-R modified HDR donor templates were used and cells were incubated with Alt-R HDR Enhancer V1 (Figure 6C). The donor templates targeting *HPRT1* had 5.9% HDR using an unmodified DNA donor template, which was increased 4.8-fold to 28.4% using an Alt-R modified donor template. The addition of Alt-R HDR Enhancer V1 further increased the frequency of HDR 1.8-fold to 52.0%. Similarly, HDR at the *MYC* locus increased from 15% HDR using an unmodified donor template to 57.5% HDR with the combined use of an Alt-R modified donor template and Alt-R HDR Enhancer V1. The same was true for *SAA1*, which had HDR frequency of 4.1% with an unmodified donor template, 18.5% with an Alt-R modified donor template, and 39.4% HDR with the combination of Alt-R modified donor template and Alt-R HDR Enhancer V1, an overall increase of 9.6-fold.

## DISCUSSION

ssODN donor templates are routinely used to generate mutations or small insertions with CRISPR-Cas proteins. This is desirable for many applications including the generation of functional domains such as epitope tags or fluorescent proteins fused to endogenous genes for biological studies, creation of cell lines with a known mutation for disease modeling, and correction of a genetic disease for therapeutic applications.^36, 51, 52^ However, the design of these donor templates remains challenging for researchers due to uncertainty about which CRISPR-Cas system should be applied, selection of gRNA(s) for each new HDR mutation location, and which donor template strand should be used to achieve the highest frequency of HDR. In addition, the design process for ssODN donors can be time-consuming, particularly if the researcher wishes to add silent blocking mutations to prevent re-cleavage and maintain amino acid translation. We have thoroughly investigated design considerations for *S.p.* Cas9 nuclease, *S.p.* Cas9 D10A nickase and *A.s.* Cas12a nuclease and present optimized design considerations for each enzyme, including positioning of the gRNA(s) relative to the desired mutation, donor strand preference, and the incorporation of blocking mutations to improve desired HDR. Additionally, we have identified donor template chemical modifications and small molecule compounds that further increase rates of HDR.

When starting an HDR genome editing project the first consideration is which CRISPR-Cas enzyme to utilize. Our results support that this choice should be dependent on where the relative genomic location of the desired mutation(s) resides in relation to the available CRISPR-Cas guides. If there is an ‘NGG’ PAM near the desired mutation (<15 bases), and this guide is expected or known to edit efficiently, then WT Cas9 can be used with confidence. If the available ‘NGG’ PAM sites are greater than 15 bases from the desired mutation, then the use of a PAM-out paired guide design with Cas9 D10A nickase may confer higher HDR than WT Cas9, provided the mutation is placed between the two nick sites generated by Cas9 D10A. This can be particularly useful if additional blocking mutations are not desired or off-target DSBs are a concern. Alternatively, if there is a ‘TTTV’ PAM site that is positioned so the HDR mutation lies between the 12-16^th^ bases of a Cas12a protospacer, then Cas12a is a viable option, although, like *S.p.* Cas9, this window can be extended with the incorporation of blocking mutations. When multiple gRNA options are available for a desired HDR edit, screening several may help eliminate any low activity guides to determine which will yield the highest HDR.

The availability of efficient Cas9 guides near a desired mutation is a significant limitation for many HDR experiments. In many cases, the guide or guides closest to the desired HDR mutation are sub-optimal in terms of cleavage efficiency or proximity. While using a paired-guide nickase strategy is viable, the requirement of having two guides with optimal spacing, activity, and orientation limits the design options and precludes this strategy for certain sites where there are no nickase designs available. Paix et al.^44^ demonstrated that incorporating additional mutations in the repair track between the cut site and desired HDR mutation location facilitated a wider region of donor integration. We observed this to be beneficial for PAM-distal HDR events. However, repair track mutations in combination with PAM mutations did not yield the highest HDR. This is likely because the repair track mutations were sufficient to prevent Cas9 re-cleavage without additional mutations in the PAM and the PAM mutation is on the opposite side of the Cas9 cleavage event, which may fall outside of the effective conversion zone for SDSA repair.^38^ For PAM-proximal HDR mutations, repair track mutations were beneficial for some sites, but not all, indicating that this strategy is effective in certain cases. In situations where optimally spaced Cas9 or Cas12a guide designs are not possible, incorporating mutations within the repair track between the cut site and desired HDR mutation or within the PAM may improve the rate of successful HDR.

The addition of blocking mutations has been demonstrated to improve HDR depending on the selected guide RNA and its relative positioning to the desired HDR mutation. Blocking mutations are beneficial when the desired HDR mutation does not prevent re-cleavage by the CRISPR-Cas nuclease. We have optimized the design of blocking mutations for use with Cas9 nuclease, including the placement and number of blocking mutations required, and this has been built into the Alt-R HDR Design Tool which facilitates simple donor template design in an easy to navigate interface. The Alt-R HDR Design Tool allows for gRNA selection for both WT Cas9, balancing the distance from the cut to mutation and on- and off-target scores of available gRNAs, and Cas9 D10A nickase, where the gRNA orientation and distance between nick sites is considered. In addition, the Alt-R HDR Design Tool provides the option to add silent blocking mutations using our empirically defined ruleset. In our study, we identified no bias in which alternate base was used as the blocking mutation to prevent Cas9 re-cleavage, indicating that there is flexibility in designing appropriate silent blocking mutations so as to not affect coding sequence. However, this could also indicate that the effect size is small or site-specific and with a larger data set potential differences between alternate bases used for silent blocking mutations could be resolved. Further investigation into the optimized number and placement of blocking mutations with Cas12a is underway with the expectation that this will be built into a tool for Cas12a HDR donor template design.

After a CRISPR-Cas system and gRNA(s) have been selected and the donor template has been designed, the next consideration for HDR experiments is the selection of homology arm lengths. In previous work investigating HDR improvements in the cell lines mentioned above, asymmetric homology arms did not improve HDR beyond symmetric homology arms when arm length was ≥30-nt from both the mutation location and the Cas9 cleavage site (data not shown). As such, the standard approach we employ is to design ssODN donor templates with 40-nt homology arms. The Alt-R HDR Design Tool allows for custom homology arm lengths to accommodate asymmetric designs, if desired. A final donor template design consideration that we investigated was strand preference for the donor template. Cas9 D10A nickase did not demonstrate a strong strand preference, so testing both strands to determine which results in the highest HDR frequency may be prudent. However, for WT Cas9 the preferred strand is strongly dependent upon where the desired HDR mutation is, relative to the Cas9 gRNA. Previous reports have demonstrated that when using ssODN donor templates with Cas9 nuclease the SDSA mechanism of repair is preferentially utilized, which consists of two steps.^38^ After a DSB is generated, the ends are resected, generating 3’ overhangs which are then available for base pairing with the donor DNA. This donor DNA then serves as a template for 5’ to 3’ DNA synthesis. Although we observed no universal donor strand preference in the experiment outlined in Figure 1B, the HDR insertion was placed directly at the Cas9 cleavage site where the SDSA model predicts high relative HDR regardless of the donor strand used. However, for insertions further from the Cas9 cleavage site there is a preference for the donor strand that contains 3’ sequence complementary to the overhangs generated during DSB repair.^44^ For PAM-proximal insertions the T strand should be used, and for PAM-distal mutations the NT strand should be used, consistent with the SDSA model of DSB repair using ssODN donor templates. For mutations directly at the cut site, we provide some evidence that the use of the T strand may reduce total editing with Cas9 which negatively impacts HDR, but this was not the case for both cell types tested. Using Cas12a, we observed a reduction in total editing rates when the T strand was used universally. We hypothesize that the donor template acts as a sponge for RNP, reducing the concentration available for genome editing within cells, or activates the non-specific ssDNase activity of Cas12a. The NT strand conferred increased HDR for experiments with Cas12a over the T strand. However, the effect of HDR insertion placement has not been thoroughly investigated for Cas12a to determine if the T strand will be advantageous over the NT strand for PAM-proximal mutations in a manner similar to Cas9 and further experimentation is required.

Beyond donor template design considerations, the use of optimized reagents for HDR experiments can further improve the frequency of HDR-mediated repair. The use of end-protecting modifications to stabilize ssODN donor templates within the cellular environment has been demonstrated to improve HDR rates, and we have developed a novel end-blocking modification that confers the highest level of improvement over unmodified DNA templates, compared to previously reported constructs. Further, we have identified two small molecules that can be used to inhibit the NHEJ pathway to improve HDR over untreated cells. We have demonstrated that when incorporated into our workflows these compounds improve HDR up to 3.2-fold in immortalized cell lines and function in combination with modified donor templates to provide further improvements in HDR frequencies. We have studied design rules for *A.s.* Cas12a nuclease, which had not yet been systematically examined. Further, the ruleset for *S.p*. Cas9 and *S.p*. Cas9 D10A nickase have been incorporated into a novel bioinformatic tool for HDR donor template design. Taken altogether, these findings present design recommendations and optimized reagents for achieving high frequency of precise repair outcomes utilizing HDR in mammalian cell lines.

## METHODS

### Ribonucleoprotein complex formation

Cas9 gRNAs were prepared by mixing equimolar amounts of Alt-R™ crRNA and Alt-R tracrRNA (Integrated DNA Technologies, Coralville, IA, USA) in IDT Duplex Buffer (30 mM HEPES, pH 7.5, 100 mM potassium acetate; Integrated DNA Technologies), heating to 95°C and slowly cooling to room temperature or using Alt-R sgRNA (Integrated DNA Technologies) hydrated in IDTE pH 7.5 (10 mM Tris, pH 7.5, 0.1 mM EDTA; Integrated DNA Technologies). Cas12a gRNAs consisted of Alt-R Cas12a crRNAs (Integrated DNA Technologies) hydrated in IDTE pH 7.5. RNP complexes were assembled by combining the CRISPR-Cas nuclease (Alt-R *S.p.* Cas9 Nuclease V3, Alt-R *S.p.* HiFi Cas9 Nuclease V3, Alt-R *S.p.* Cas9 D10A V3, Alt-R *S.p.* Cas9 H840A V3, Alt-R *A.s.* Cas12a V3, or Alt-R *A.s.* Cas12a *Ultra*; Integrated DNA Technologies) and the Alt-R gRNA at a 1:1 to 1.2:1 molar ratio of gRNA:protein and incubating at room temperature for 30 minutes. For paired nicking experiments, each RNP was formed separately, and two RNPs were mixed together at an equal molar ratio prior to adding to the cells at the time of transfection. The 20-nt target specific sequences of the gRNAs used in this study are listed in Supplementary Table 1.

### HDR ssODN donor templates

Alt-R™ HDR Donor Oligos (Integrated DNA Technologies) used in this study consisted of either Alt-R modified (containing two phosphorothioate linkages at the ultimate and penultimate backbone linkage and an IDT proprietary end-blocking modification at 5’ and 3’ ends), PS modified (containing two phosphorothioate linkages at the ultimate and penultimate backbone linkage at 5’ and 3’ ends) or unmodified DNA. Donor oligos were hydrated using IDTE pH 7.5 (Integrated DNA Technologies). Sequences of the HDR oligos used in this study are listed in Supplementary Table 1.

### Cell culture

HAP1, HEK293, HeLa, Jurkat E6-1, and K562 cells were purchased from ATCC^®^ (Manassas, VA, USA), and maintained in DMEM (HEK293, and HeLa), RPMI-1640 (Jurkat) and IMDM (HAP1, K562) (ATCC), each supplemented with 10% fetal bovine serum (FBS) and 1% penicillin-streptomycin (Thermo Fisher Scientific, Carlsbad, CA, USA). HEK293 cells that constitutively express Cas9 nuclease (”HEK293-Cas9”) were generated by stable integration of a human-codon optimized *S.p.* Cas9 as well as the flanking 5’ and 3’ nuclear localizing sequences and 5’-V5 tag from the GeneArt CRISPR Nuclease Vector (Thermo Fisher Scientific). HEK293-Cas9 cells were maintained in DMEM supplemented with 10% FBS, 1% penicillin-streptomycin, and 500 μg/mL G418 (Thermo Fisher Scientific). Cells were incubated at 37°C with 5% CO2 and passaged every 3 days. HAP1 cells were used for transfection at 50-70% confluency. HEK293 and HeLa cells were used for transfection at 70-90% confluency. Jurkat and K562 were used for transfection at 5-8 x 10^5^ cells/mL density. After transfection, cells were grown for 48-72 hours in total, after which genomic DNA was isolated using QuickExtract^TM^ DNA Extraction Solution (Epicentre, Madison, WI, USA).

### Delivery of genome editing reagents by lipofection

Lipofection was performed in 96-well plates. First, 25 µL of Opti-MEM^®^ (Thermo Fisher Scientific) containing 1.2 µL (RNP delivery) or 0.75 µL (gRNA delivery) of Lipofectamine^®^ RNAiMAX (Thermo Fisher Scientific) was combined with equal volume of Opti-MEM containing RNP or gRNA and HDR donor template (when present), and incubated at room temperature for 20 min. After lipoplex formation, 4.5 x 10^4^ cells resuspended in 100 µL of DMEM + 10% FBS were added to the transfection complex which resulted in a final concentration of 10 nM RNP or gRNA and 3 nM HDR oligo on a per-well basis. Transfection plates were incubated at 37°C and 5% CO2.

### Delivery of genome editing reagents by nucleofection

Electroporation was performed using the Lonza™ Nucleofector™ 96-well Shuttle™ System (Lonza, Basel, Switzerland). For each nucleofection, cells were washed with 1X phosphate buffered saline (PBS) and resuspended in 20 µL of solution SF or SE (Lonza). Cell suspensions were combined with RNP complex(es), Alt-R Cas9 or Cpf1 (Cas12a) Electroporation Enhancer (Integrated DNA Technologies) and HDR donor template (if applicable). This mixture was transferred into one well of a Nucleocuvette™ Plate (Lonza) and electroporated using manufacturer’s recommended protocols (except for HEK293, which used protocol 96-DS-150). After nucleofection, 75 µL pre-warmed culture media was added to the cell mixture in the cuvette, mixed by pipetting, and 25 µL was transferred to a 96-well culture plate with 175 µL pre-warmed culture media. Transfection plates were incubated at 37°C and 5% CO2.

### Addition of Alt-R HDR Enhancer

For experiments using HDR Enhancer, cells were transfected as described. Immediately following transfection, cells were grown in media containing either DMSO as a vehicle control, Alt-R™ HDR Enhancer V1 at a final concentration of 30 µM, or Alt-R HDR Enhancer V2 at a final concentration of 1 µM. 24 hours after transfection, media was aspirated away without disturbing the cells and fresh media was added to each well.

### T7 Endonuclease I (T7EI) Assay and restriction enzyme digestion

Genomic DNA was extracted after 48-72 hrs incubation using 50 µL Quick Extract™ DNA Extraction Solution (Lucigen, Middleton, WI, USA) following the manufacturer’s protocol. Genomic DNA was diluted 3-fold with nuclease-free water and 1.5 µL was PCR-amplified using 0.15 U KAPA HiFi HotStart DNA Polymerase (Kapa Biosystems, Wilmington, MA, USA) in a final volume of 10 µL. For HDR analysis using restriction enzyme digestion, 10 µL of the PCR product was incubated with 2 U of EcoRI-HF^®^ in 1X CutSmart^®^ Buffer (New England BioLabs, Ipswich, MA, USA) at 37°C for 60 minutes. Total editing rate was measured using the Alt-R™ Genome Editing Detection Kit (T7EI) (Integrated DNA Technologies) following the manufacturer’s protocol. Cleavage products were separated on the Fragment Analyzer™ using the CRISPR Mutation Discovery Kit (Agilent Technologies, Santa Clara, CA, USA). Editing and HDR frequencies were calculated using the following formula: average molar concentration of the cut products / (average molar concentration of the cut products + molar concentration of the uncut product) x 100. PCR primers are listed in Supplementary Table 1.

### Quantification of editing events by next-generation sequencing (NGS)

On-target editing and HDR efficiencies were also measured by NGS. Libraries were prepared using an amplification-based method as described previously ^53^. In short, the first round of PCR was performed using target specific primers, and the second round of PCR incorporates P5 and P7 Illumina adapters to the ends of the amplicons for universal amplification. Libraries were purified using Agencourt^®^ AMPure^®^ XP system (Beckman Coulter, Brea, CA, USA), and quantified with qPCR before loading onto the Illumina^®^ MiSeq platform (Illumina, San Diego, CA, USA). Paired end, 150 bp reads were sequenced using V2 chemistry. Data were analyzed using a custom-built pipeline. Data was demultiplexed using Picard tools v2.9 (https://github.com/broadinstitute/picard). Forward and reverse reads were merged into extended amplicons (flash v1.2.11)^54^ before being aligned against the GRCh38 genomic reference (minimap2 v2.12).^55^ Reads were aligned to the target, favoring alignment choices with indels near the predicted cut site(s). At each target, editing was calculated as the percentage of total reads containing an indel within an 8bp window of the cut site for Cas9 or a 9bp window from the -3 position of the Cas12a PAM distal cut site. PCR primers are listed in Supplementary Table 1.

### Statistical Analysis

The data collected from experiments were analysed on Graph PadPrism 8 using two-tailed unpaired t-test to evaluate significance (*p < 0.05, **p < 0.01, ***p < 0.001, and ****p < 0.0001).

## DATA AVAILABILITY

The Alt-R HDR Design Tool is a free online tool that is available from the Integrated DNA Technologies website (https://www.idtdna.com/pages/tools/alt-r-crispr-hdr-design-tool). NGS data used for the figures and supplementary figures have been made available at SRA BioProject Accession # PRJNA638623.

## Supporting information

Supplemental Sequences

## ACKNOWLEDGEMENTS

The authors thank Ashley Jacobi and Kim Lennox for editing the manuscript. This work was supported by internal funds from Integrated DNA Technologies, Inc. The authors are employed by Integrated DNA Technologies, Inc., (IDT) which offers reagents for sale similar to some of the compounds described in the manuscript. GRR owns equity in DHR, the parent company of IDT.

## AUTHOR CONTRIBUTIONS

Conceptualization, M.S.S, B.T., J.W., R.T., S.Y., M.S.M. and G.R.R; Software, G.K. and M.S.M.; Investigation, M.S.S., B.T., J.W., R.T. and S.Y.; Data Curation, M.S.S. and M.S.M.; Writing-Original Draft, M.S.S.; Writing-Review & Editing, M.S.S., B.T., J.W., R.T., S.Y., G.K., M.S.M. and G.R.R.; Supervision: G.R.R.

---

**Supplemental Figure 1.**
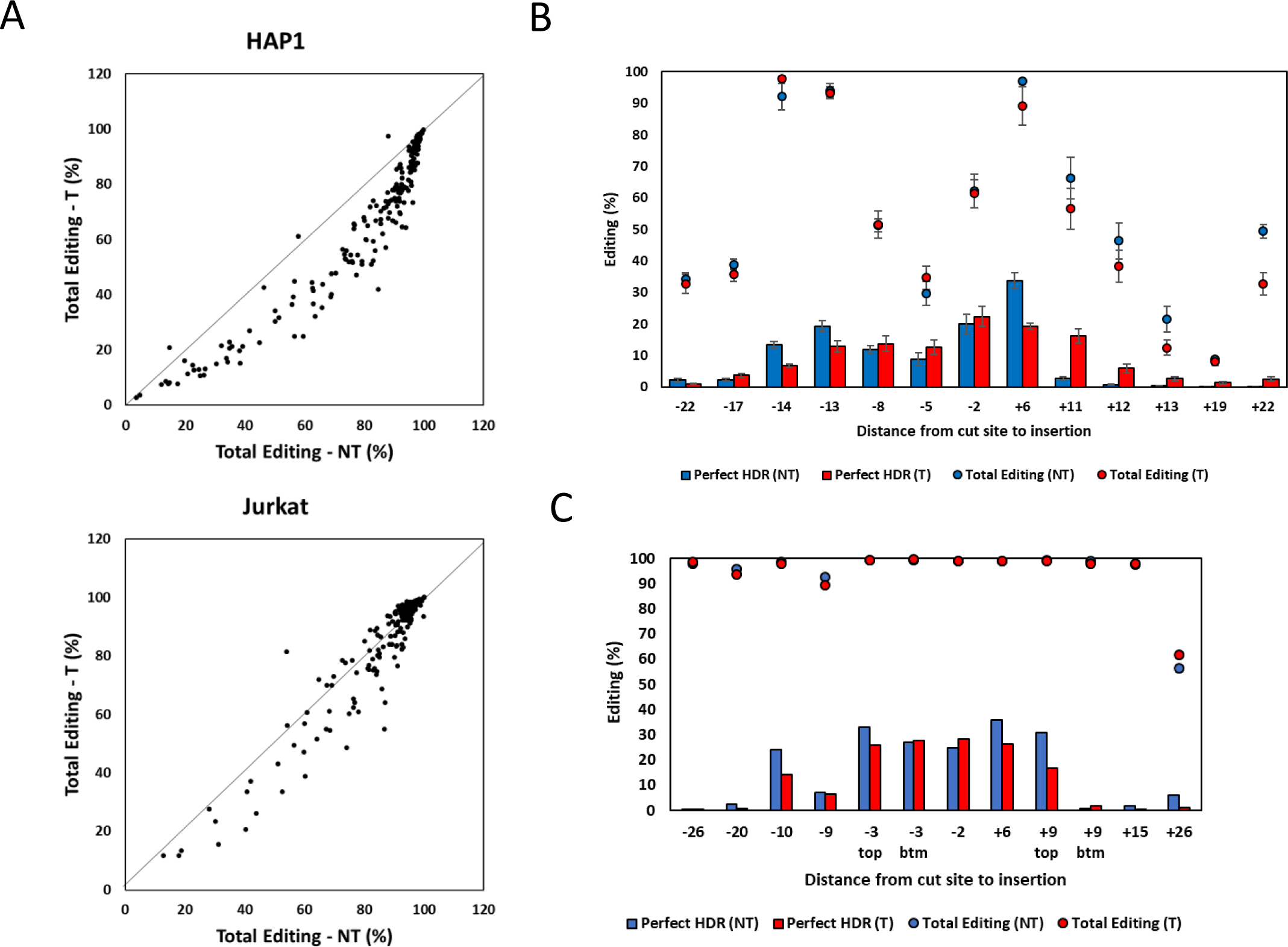
(A) EcoRI restriction digest recognition site (GAATTC) was inserted at the Cas9 cleavage site of 254 genomic loci in Jurkat and 239 genomic loci in HAP1 cells using either the targeting (T) or non-targeting (NT) strand as the donor template. RNP complexes (Alt-R *S.p.* Cas9 Nuclease complexed with Alt-R CRISPR-Cas9 crRNA and tracrRNA) were delivered at 4 µM along with 4 µM Alt-R Cas9 Electroporation Enhancer and 3 µM donor template by nucleofection. Total editing was assessed via NGS. (B) Insertion of an EcoRI site before the stop codon of GAPDH in HEK293 cells using guides around the desired HDR insertion location. The cleavage sites and associated distance to the desired insertion location for each guide are indicated above the sequence shown. Both the T and NT strand were tested. RNP complexes (Alt-R *S.p.* Cas9 Nuclease complexed with Alt-R CRISPR-Cas9 crRNA and tracrRNA) were delivered at 2 µM along with 2 µM Alt-R Cas9 Electroporation Enhancer and 2 µM donor template by nucleofection. HDR and total editing were assessed via NGS. Data are represented as means ± S.E.M. of three technical replicates. (C) Insertion of an EcoRI site at the TNPO3 locus in HEK293 cells using guides around the desired HDR insertion location. The distance from each cleavage site to the desired insertion location for each guide are indicated on the x-axis. Two pairs of guides cut at the same location, but on opposite strands. The strand containing the guide is indicated as top or bottom (btm). Both the T and NT strand were tested. RNP complexes (Alt-R *S.p.* Cas9 Nuclease complexed with Alt-R CRISPR-Cas9 crRNA and tracrRNA) were delivered at 2 µM along with 2 µM Alt-R Cas9 Electroporation Enhancer and 2 µM donor template by nucleofection. HDR and total editing were assessed via NGS.

**Supplemental Figure 2.**
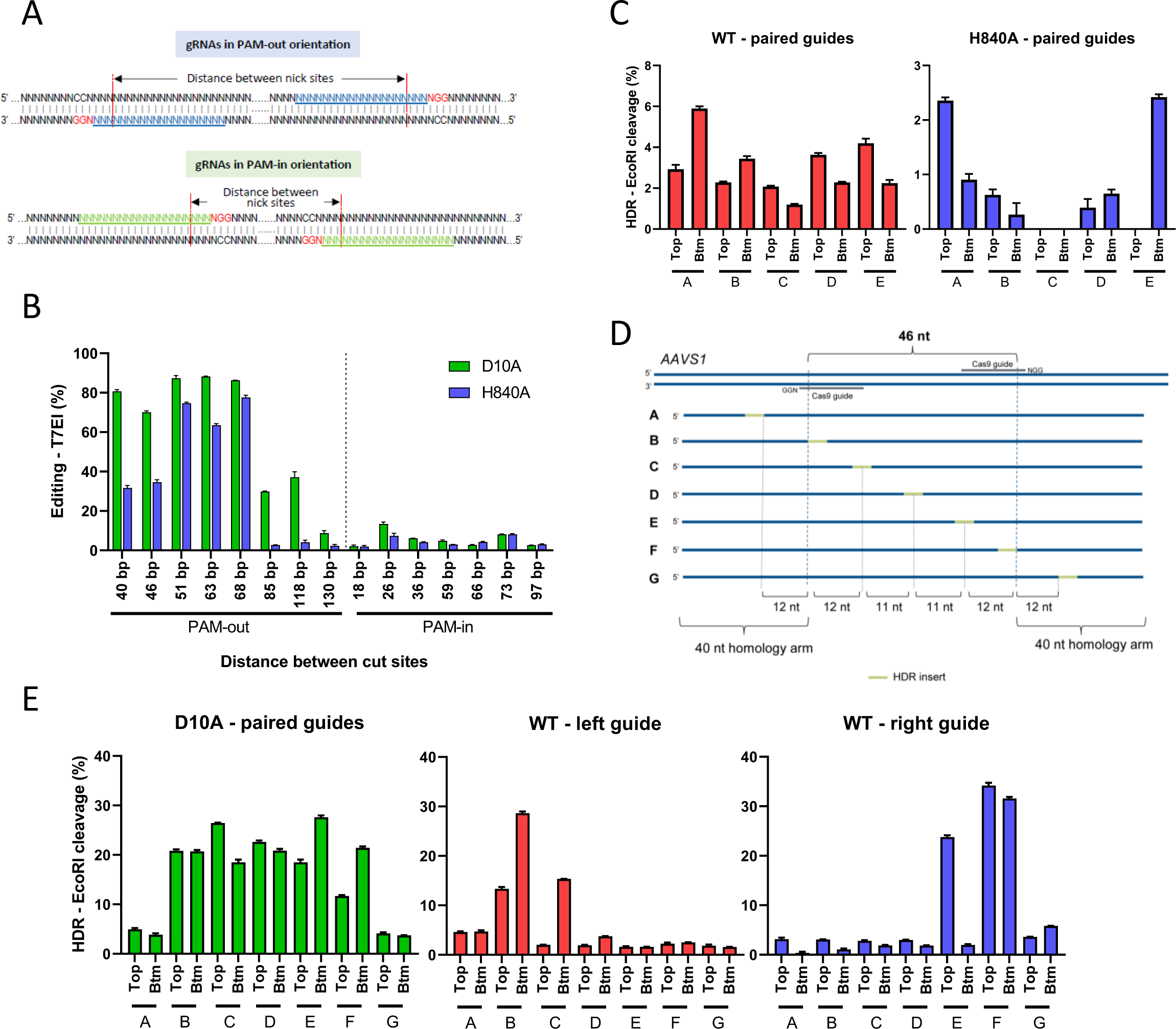
HDR using Cas9 D10A nickase compared to WT Cas9 and Cas9 H840A nickase (A) Schematics showing gRNA pairs in PAM-out orientation (top panel) or PAM-in orientation (bottom panel). NGG PAMs are red, protospacers are underlined. Spacing between paired gRNAs is defined by the distance between targeted nick sites as indicated in the diagram. (B) RNP complexes consisting of gRNA pairs in different orientation and spacing targeting the *HPRT1* locus were delivered into HEK293 cells with Cas9 D10A or H840A proteins via lipofection and total editing was measured by T7EI cleavage. (C) HDR mediated by Cas9 WT (left panel) or Cas9 H840A (right panel) with paired gRNAs. Cas9 WT and Cas9 H840A were used in combination with gRNA pairs targeting *HPRT1* 51-nt PAM-out site. RNP complexes (Alt-R *S.p.* Cas9 Nuclease or Alt-R *S.p.* Cas9 H840A nickase complexed with Alt-R CRISPR-Cas9 crRNA and tracrRNA) were delivered at 4 µM (2 µM each RNP for nickase paired guides) along with 4 µM Alt-R Cas9 Electroporation Enhancer and 2 µM donor template by nucleofection. The same set of ssODNs homologous to either the top or bottom (Btm) strand as shown in Figure 2 were used to insert an EcoRI site along the target region. HDR was assessed via EcoRI cleavage. (D) Schematics of HDR donor oligos. HDR donor sequences were designed to insert an EcoRI site at 7 positions along the AAVS1 46-nt PAM-out target region. (E) HDR performance of donor oligos in HEK293 cells. Cas9 D10A with two guides, or Cas9 WT with each of the individual guides were used to induce double strand breaks. Bar charts are showing the HDR rate using indicated oligos homologous to either the top or bottom strand. Data are represented as means ± S.E.M of technical triplicates.

**Supplemental Figure 3.**
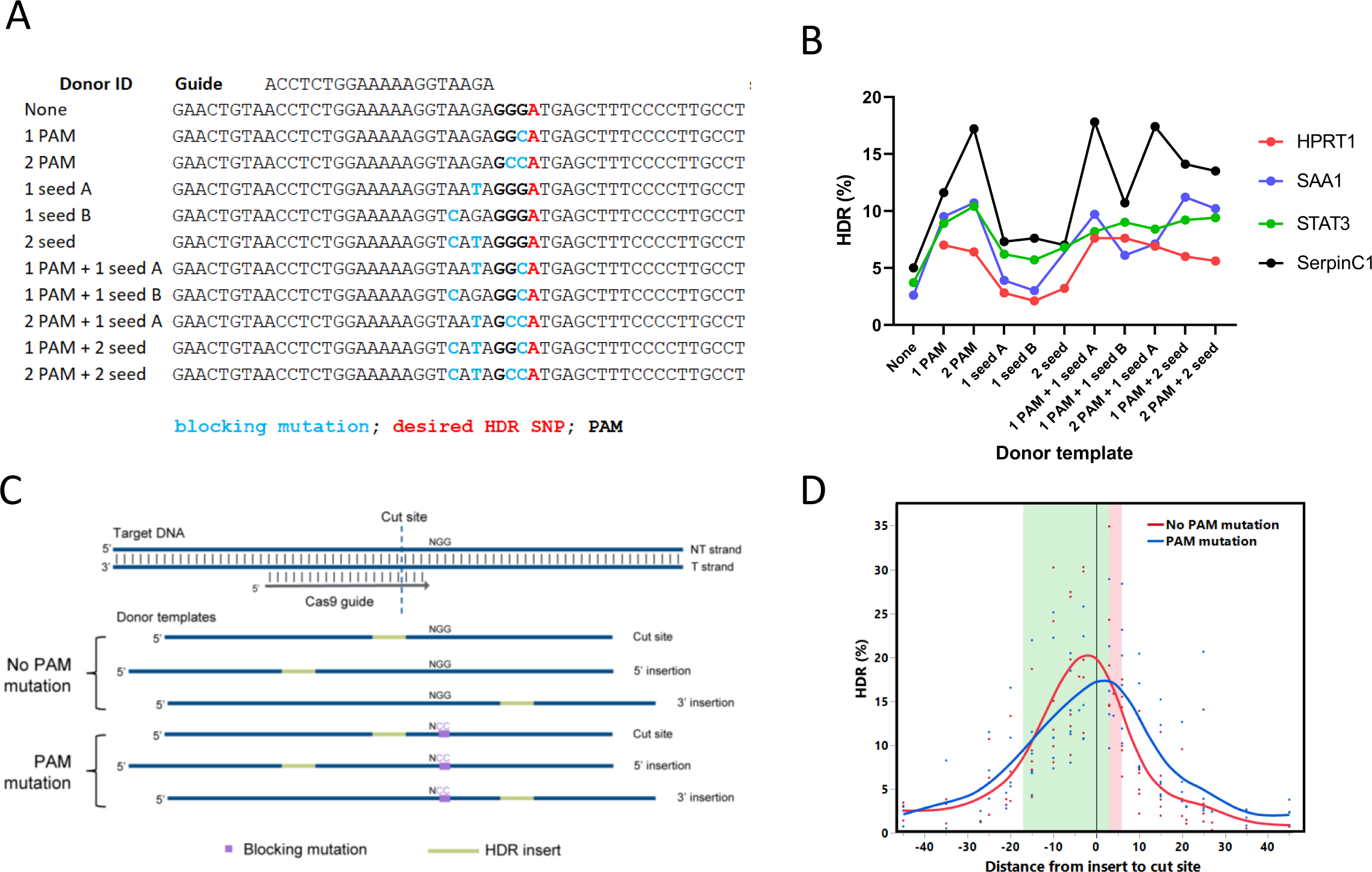
(A) Example sequence of site SERPINC1 showing the various blocking mutation(s) tested. The Cas9 guide is shown above the sequence, and the PAM is bolded. The intended HDR mutation is in red and blocking mutations are shown in blue. (B) Donor templates for four target loci following the design strategy shown in panel A were delivered to Jurkat cells at 4 µM along with RNP complexes (Alt-R *S.p.* Cas9 Nuclease complexed with Alt-R CRISPR-Cas9 crRNA and tracrRNA) at 4 µM and with 4 µM Alt-R Cas9 Electroporation Enhancer by nucleofection. SNP conversion of the desired HDR mutation 3’ of the PAM was determined by NGS. (C) Schematic representation of donor templates used to an EcoRI insert sequence positioned at varying distances from the Cas9 cleavage site, ranging up to 45 bases in either the 5’ or 3’ direction. Donor templates were designed with and without a mutation in the PAM (‘NGG’ to ‘NCC’) to prevent Cas9 re-cleavage. (D) HDR performance of donor templates for four genomic loci in Jurkat cells and two genomic loci in HEK293 cells. Negative values indicate the insertion was 5’ (PAM-distal) of the cut site, whereas positive values indicate the insertion was 3’ (PAM-proximal) of the cut site. RNP complexes (Alt-R *S.p*. Cas9 Nuclease complexed with Alt-R CRISPR-Cas9 crRNA and tracrRNA) were delivered at 4 µM along with 4 µM Alt-R Cas9 Electroporation Enhancer and 4 µM donor template by nucleofection. HDR rates were assessed via EcoRI cleavage of targeted amplicons.

**Supplemental Figure 4.**
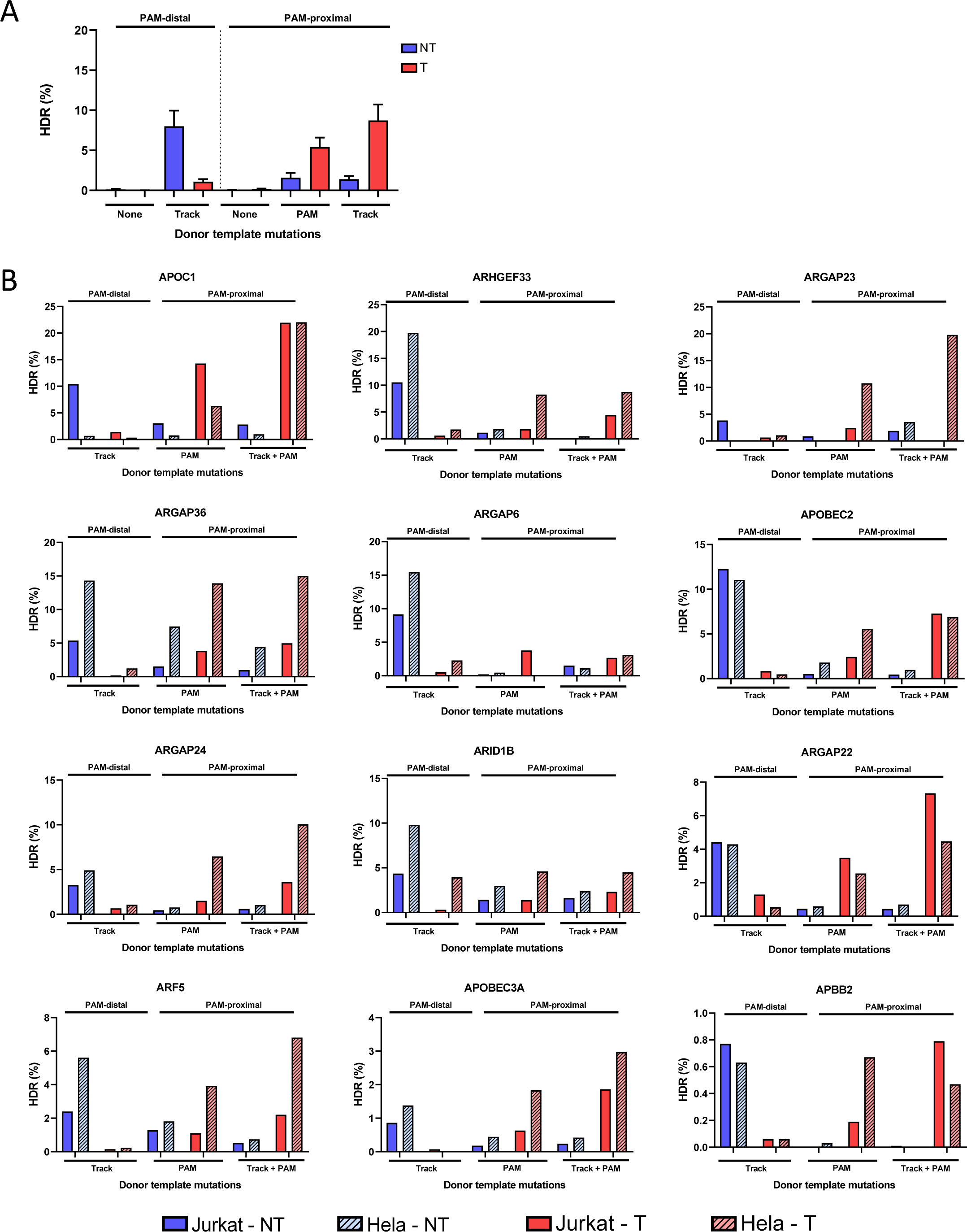
(A) Donor templates creating an EcoRI insertion at the cut site or 20 bases PAM-proximal or PAM-distal to the Cas9 cut site for 12 genomic loci were tested in Hela cells as the targeting (T) or non-targeting (NT) strand. Donor templates for the PAM-distal insert contained repair track mutations. Donor templates for the PAM-proximal insert contained either PAM mutation or repair track plus PAM mutations. RNP complexes (Alt-R *S.p.* Cas9 Nuclease complexed with Alt-R CRISPR-Cas9 sgRNA) were delivered at 4 µM along with 4 µM Alt-R Cas9 Electroporation Enhancer and 3 µM donor template by nucleofection. Perfect HDR rates were determined by NGS. Data are represented as means ± S.E.M. (B) Individual plots of the 12 sites tested in HeLa and Jurkat cells.

**Supplemental Figure 5.**
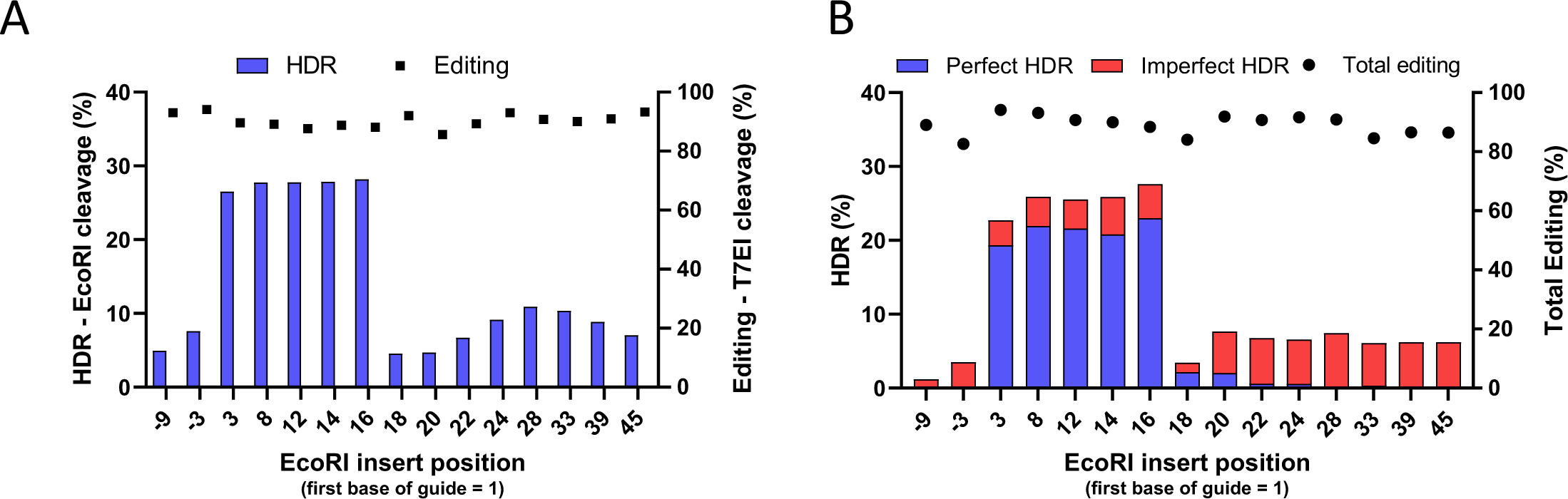
Site HPRT 38330 from Figure 5B was delivered as an RNP complex (Alt-R *A.s.* Cas12a *Ultra* nuclease complexed with Alt-R CRISPR-Cas12a crRNA) at 2 µM along with 3 µM Alt-R Cpf1 Electroporation Enhancer and 3 µM Alt-R modified donor templates by nucleofection to Jurkat cells. HDR was measured by EcoRI cleavage (A) and NGS analysis (B) to determine the frequency of perfect HDR (blue) relative to imperfect HDR (red) and total editing (black dots).

